# Spinal dI2 interneurons regulate the stability of bipedal stepping

**DOI:** 10.1101/2020.01.07.898072

**Authors:** Baruch Haimson, Yoav Hadas, Artur Kania, Monica Daley, Yuval Cinnamon, Aharon Lev-Tov, Avihu Klar

## Abstract

Peripheral and intraspinal feedback is required to shape and update the output of spinal networks that execute motor behavior. We report that lumbar dI2 spinal interneurons of the chick receive synaptic input from afferents and pre-motoneurons. They innervate contralateral premotor networks in the lumbar and brachial spinal cord and their ascending projections innervate the cerebellum. These findings suggest that dI2 neurons function as interneurons in local lumbar circuits and involved in lumbo-brachial coupling and that part of them deliver peripheral and intraspinal feedback to the cerebellum. Silencing of dI2 neurons leads to destabilized stepping in P8 hatchlings with occasional collapses, variable step-profiles and wide-base walking, suggesting that the dI2 neurons may contribute to stabilization of the bipedal gait.

## Introduction

The spinal cord integrates and relays the somatosensory inputs required for further execution of complex motor behaviors. Interneurons that differentiate at the ventral progenitor domain, V3-V0, are involved in the control of rhythmic motor activity, alternating between left and right limbs, as well as between flexor and extensor muscles (Lai et al., 2016, Osseward and Pfaff, 2019, Alaynick et al., 2011). Some of the dorsally born interneurons, dI1, dI3 and dI6, migrate ventrally and are also assembled within circuitry that control motor activity (Yuengert et al., 2015, Bui et al., 2013, Andersson et al., 2012), while other dorsal progenitor neurons, dI4 and dI5, give rise to interneurons that mediate somatosensation (Lai et al., 2016). The role of dI2 neuron is elusive due to the lack of genetic targeting means. Employing intersectional genetics in the chick spinal cord we targeted dI2 and present evidences that implicate them in the control of stability during locomotion.

The maintenance of stability of the body and the coordination, precision and timing of movements are regulated and modulated by the cerebellum. Anatomical and electrophysiological studies of cats and rodents revealed two major pathways ascending from tract neurons in the lumbar spinal cord to the cerebellum: the dorsal and the ventral spinocerebellar tract (DSCT and VSCT). DSCT tract neurons are considered to relay mainly proprioceptive information, while VSCT tract neurons are thought to relay internal spinal network-information to the cerebellum in addition to proprioceptive data (Jankowska and Hammar, 2013, Spanne and Jorntell, 2013, Stecina et al., 2013, Jiang et al., 2015). While subpopulations of DSCT neuron are genetically accessible (Hantman and Jessell, 2010), the genetic inaccessibility of VSCT neurons hinders revealing their actual contribution to the regulatory functions of the cerebellum in locomotion and other motor behaviors.

In the present study we studied the possible functions of dI2 neurons in chick motor behavior. There are several reasons and advantages to perform these studies in chicks: The patterning of neurons within the spinal cord (Jessell, 2000) and the spinocerebellar tracts (Furue et al., 2010, Furue et al., 2011, Uehara et al., 2012) are conserved between mammalians and avian. In addition, chick uses bipedal locomotion (evolved in humans and birds) that can be examined soon after hatching. For decoding the circuitry and function of spinal interneurons we developed a unique circuit-deciphering toolbox that enables neuronal-specific targeting and tracing of circuits in the chick embryo (Hadas et al., 2014), and utilized kinematic analysis of overground bipedal stepping of the hatched chicks following silencing of dI2.

Our studies revealed that lumbar dl2 neurons receive synaptic inputs from inhibitory and excitatory premotoneurons and relay output to the cerebellar granular layer, premotor neurons at the contralateral spinal cord and the contralateral dI2. Kinematic analysis of overground stepping of P8 hatchlings after Inhibition of the neuronal activity of dI2s by targeted expression of the tetanus toxin gene, showed unstable stepping in the genetically manipulated hatchlings, hence demonstrating that dI2s play a role in shaping and stabilizing the bipedal gait.

## Results

In order to define potential VSCT neurons within spinal interneurons, we set the following criteria: 1) Soma location in accordance with pre-cerebellar neurons at the lumbar level, which were previously revealed by retrograde labeling experiments of the chick cerebellar lobes (Furue et al., 2010, Furue et al., 2011, Uehara et al., 2012), 2) Commissural neurons, 3) excitatory neurons, and 4) non-premotoneurons (Lai et al., 2016, Osseward and Pfaff, 2019, Alaynick et al., 2011). Based on these criteria dI1c and dI2 neurons are likely candidates (Bermingham et al., 2001, Yuengert et al., 2015) (Fig. S1B). This is further supported by Sakai et al., who demonstrated that in the E12 chick, dI1 and dI2 axons project to the hindbrain and toward the cerebellum (Sakai et al., 2012). In the current study we have focused on deciphering the circuitry and function of dI2 neurons and their possible association to VSCT.

### dI2 interneurons are mainly excitatory neurons with commissural axonal projection

The dI2 neurons originate at the dorsal spinal cord. The progenitor pdI2 express Ngn1, Ngn2, Olig3 and Pax3 transcription factors (TFs). The post mitotic dI2 are defined by a combinatorial expression of Foxd3^+^/Lhx1^+^/Pou4f1^+^ TFs (Alaynick et al., 2011, Morikawa et al., 2009, Francius et al., 2013). To label dI2 neurons, axons, and terminals in the chick spinal cord we used intersection between enhancers of two dI2’s TFs - Ngn1 and the Foxd3, via expression of two recombinases (Cre and FLPo) and double conditional reporters (Fig. S1A). We have shown previously that the combination of these enhancers reliably labels dI2 neurons (Avraham et al., 2009, Hadas et al., 2014). At E5, early post mitotic dI2 neurons migrate ventrally from the dorso-lateral to the mid-lateral spinal cord (Fig. 1A). As they migrate ventrally, at E6, dI2 neurons assume a mid-lateral position along the dorso-ventral axis (Fig. 1B). Subsequently, dI2 neurons migrate medially, and at E17, comparable to pos-gestation day 4 (P4) of mouse, they occupy lamina VII (Fig. 1C). At all rostro-caudal levels and embryonic stages, dI2 axons cross the floor plate (Fig. 1A-D). Post-crossing dI2 axons extend rostrally for a few segments at the ventral funiculus (VF) and subsequently turn into the lateral funiculus (LF) (Avraham et al., 2009) (Fig. 1C). The VF to LF rerouting is apparent at the rostral thoracic levels (Fig. 1D). Collaterals originating from the crossed VF and LF tracts invade the contralateral spinal cord (Fig. 1C, D, Fig. 4A,B).

**Figure 1:**
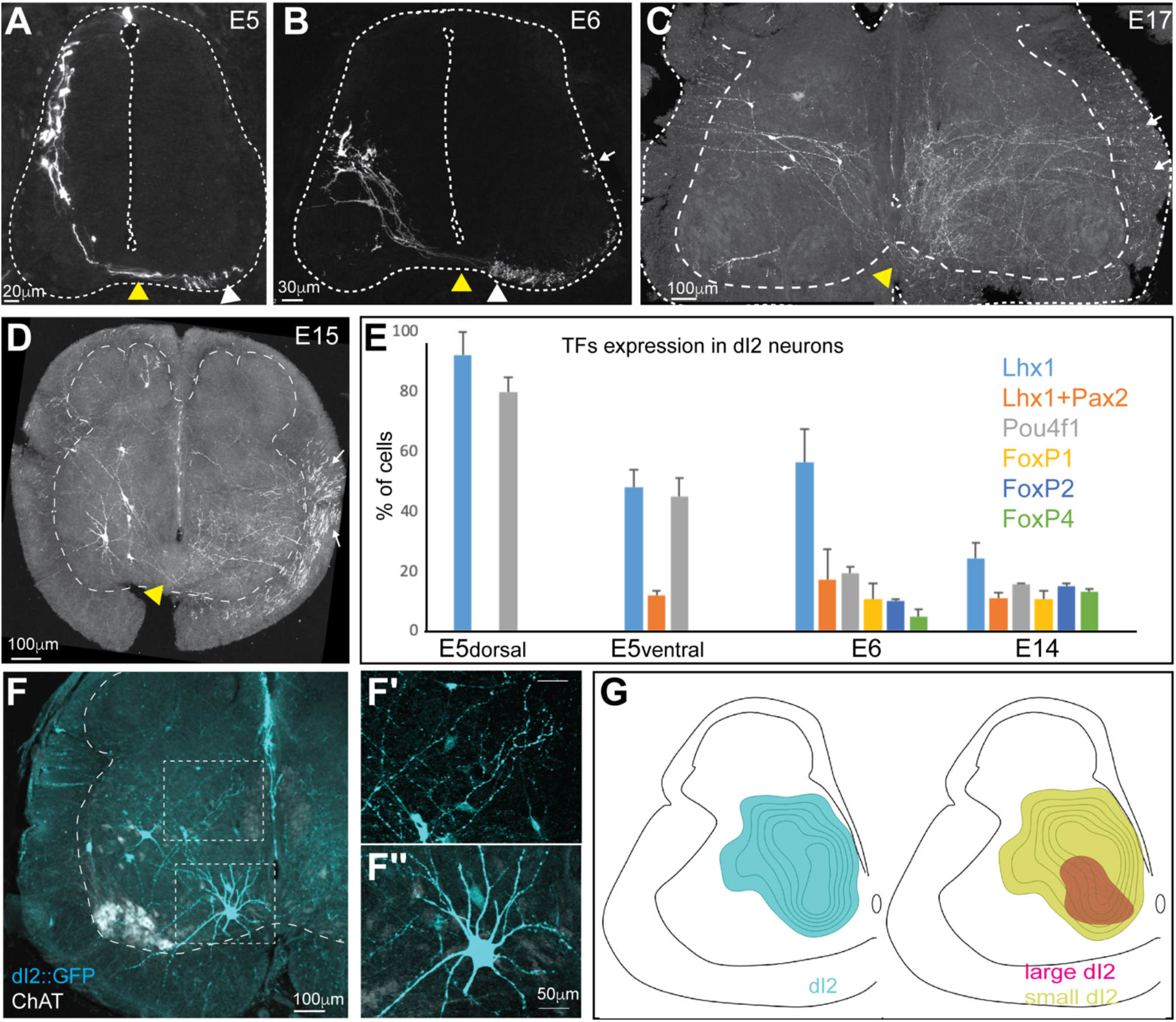
Characterization and classification of dI2 neurons during embryonic development. dI2 interneurons were labeled by genetic intersection between Foxd3 and Ngn1 enhancers (Avraham et al., 2009) (Supp Fig. S1). **A–D.** dI2 axonal projection during development. At E5 (A) post mitotic dI2 neurons assume a dorsolateral position and start to migrate ventrally. At E6 (B) dI2 neurons occupy the mid-lateral domain. At E15-17 dI2 neurons are located at the medial lamina VII at the lumbar level (LS3) (C) and the thoracic level (T1) (D). dI2 axons cross the floor plate (yellow arrowheads), turn longitudinally at the ventral funiculus (white arrowheads) and eventually elongate at the lateral funiculus (white arrows). **E.** A graph indicating the fraction of dI2 neurons expressing TFs during development (based on data from three E5, two E6 and two E14 embryos). **F.** Cross section of an E17 embryo at the lumbar spinal cord (crural plexus level, LS2). Small-diameter dI2 neurons residing in lamina VII (F’) and ventromedial large-diameter dI2 neurons in lamina VIII (F”). **G.** Density plots of dI2 somata in the sciatic plexus level (cyan, *N*=374 cells), dI2_large_ (magenta) and dI2_small_ (yellow) INs (*N*=33 and *N*=344 cells, respectively, from 2 embryos). See Figure S2 and S3.

A recent study suggested that dI2 neurons at early stages of development in mice at E9.5-E13.5 (comparable the chick E4-8) can be divided into several sub classes based on their genetic signature, and degree of maturation (Delile et al., 2019). To assess the diversity of dI2 neurons in the chick, the expression of dI2 TFs in dI2::GFP cells was analyzed at E5 before and during ventral migration, E6 and E14. The early post mitotic dI2::GFP at E5 are a homogenous population defined by Foxd3^+^/Lhx1^+^/Pou4f1^+^/Pax2^−^ (Fig. 1E, S2A,B). The dI2 neurons that undergo ventral migration at E5, as well as at E6 and E14, express variable combinations of Lhx1, Pou4f1 and FoxP1/2/4 (Fig. 1E, S2C,D). At E14, about 50% of dI2::GFP did not express any of the tested TFs (Fig. 1E), suggesting that the early expression of TFs is required for cell fate acquisition, axon guidance, and target recognition, while their expression is redundant during circuitry formation, as shown for other spinal INs (Bikoff et al., 2016). Interestingly, about 12% of ventrally migrating dI2 neurons (from E5 to E14) express Pax2 (Fig. 1E, S2D). Pax2 is associated with an inhibitory phenotype (Cheng et al., 2004), suggesting that a subpopulation of dI2 are inhibitory neurons. The distribution of excitatory and inhibitory dI2 neurons is also apparent at E17. *In situ* hybridization on cross sections of E17 dI2::GFP-labeled lumbar spinal cord, the vGlut2 probe revealed that 88% were vGlut2^+^, while the Gad1 probe measured 25% Gad1^+^ dI2 neurons (Fig. S3A-D). A similar percentage of Gad2 and Slc6a5 dI2-expressing cells was found also in mouse (Delile et al., 2019). At E13-17 in the caudal lumbar level and at the level of the sciatic plexus, most dI2 neurons reside at the medial aspect of lamina VII. About 91% of dI2 neurons are small-diameter neurons that reside at the lateral dorsal aspect of lamina VII, and 9% are large-diameter neurons. At the lumbar sciatic plexus level, large-diameter dI2 neurons mostly reside at the ventral aspect of lamina VII (Fig. 1F,G) and at the level of the crural plexus in the ventral and dorsal aspect of lamina VII (Fig. S3E). Importantly, large-diameter dI2 neurons are only apparent at the lumbar level (Fig. 1F, S3E-F). The division of large- and small-diameter lumbar dI2 neurons was not reflected in the expression of the tested TFs. Hence, dI2 neurons consist several sub-populations, as it was shown with other spinal interneurons (Bikoff et al., 2016, Delile et al., 2019, Sweeney et al., 2018)

### Sub-population of dI2 neurons project to the cerebellum

To study the supraspinal targets of dI2 neurons, axonal and synaptic reporters were expressed in lumbar dI2 neurons (Fig. 2A). At E3, dI2 enhancers were co-electroporated with double conditional axonal reporter – membrane tethered cherry, and synaptic reporter – SV2-GFP (Fig. S1A). Expression into the lumbar spinal cord was attained by using thin electrodes positioned near the lumbar segments. At E17, the stage in which the internal granule layer is formed in the chick cerebellum, the axons and synapses of dI2 neurons were studied. dI2 axons cross the spinal cord at the floor plate at the segmental level, ascend to the cerebellum, enter through the superior cerebellar peduncle, and cross back to the other side of the cerebellum ipsilaterally to the targeted dI2s (Fig. 2B). Synaptic boutons are noticeable in the granule layer at the ipsilateral and contralateral sides of the anterior cerebellar lobules (Fig. 2C). Synaptic boutons were also present in the central cerebellar nuclei (Fig. S4A).

**Figure 2:**
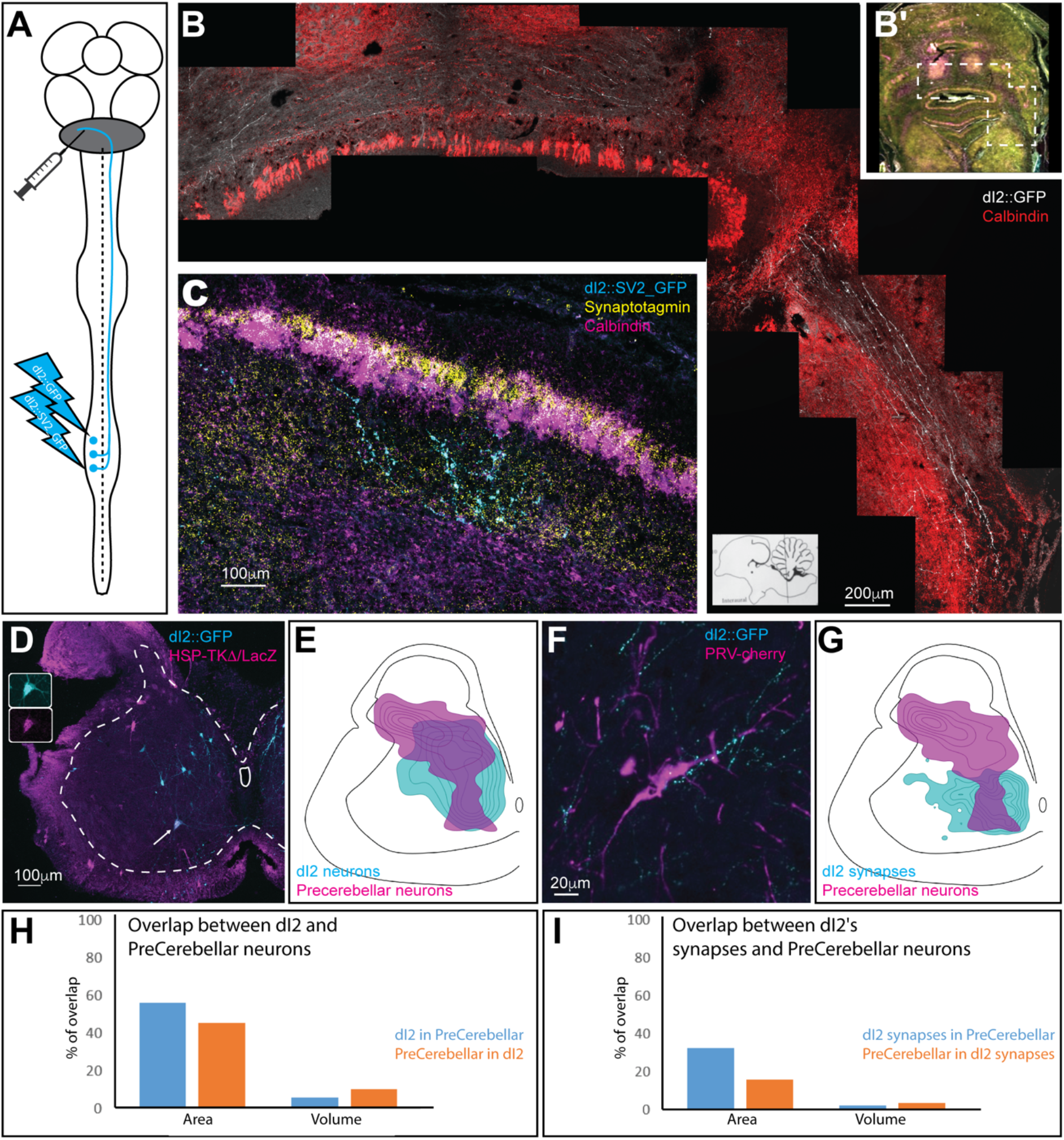
dI2 neurons project to the cerebellum. **A.** Experimental setup for labeling of cerebellar projecting dI2 neurons. dI2 neurons were genetically targeted at HH18, and pre-cerebellar neurons were labeled using intra-cerebellar injection of CTB or replication defective HSV-LacZ at E15. **B.** A cross section of E17 brainstem and cerebellum. The dashed polygon in B’ is magnified in B. dI2 axons reach the cerebellum, enter into it via the superior cerebellar peduncle and cross the cerebellum midline. Calbindin (Purkinje neurons, magenta (B’) or red (B)), synaptotagmin (yellow). **C.** A cross section of E17 cerebellar cortex. Lumbar-originating dI2 synapses (cyan) in the granular layer of the anterior cerebellar cortex. Calbindin (Purkinje neurons, magenta), synaptotagmin (yellow). **D.** A cross section of an E15 embryo at the lumbar spinal cord level (sciatic plexus level). Pre-cerebellar neurons were infected and labeled by HSV-LacZ (magenta), and dI2 neurons express GFP (cyan). A large-diameter dI2 neuron co-expressing LacZ and GFP is indicated by an arrow (magnification of the two channels in the insets). **E.** Density plots of dI2 and pre-cerebellar neurons (density values 10-90%) in the sciatic plexus segments (*N*=374 and *N*=289 cells, respectively). **F.** CTB labeled pre-cerebellar neuron (magenta) is contacted by dI2 axonal terminals (cyan). **G.** Density plots of dI2 synapses and pre-cerebellar neuron somata (density values 10-90%) in the sciatic plexus segments (*N*=4735 synapses and *N*=289 cells, respectively). **H,I.** Quantification of the overlap in area and volume of the two density plots. The plots are based on data from 3 embryos. See Figure S4.

The difference in the soma size between the dorsally and ventrally located dI2 neurons prompted us to test which dI2 neurons project to the cerebellum. The dI2 and pre-cerebellar neurons were co-labeled by genetic targeting of dI2 at early stages of embryogenesis (E3), coupled with intra-cerebellar injection of cholera toxin subunit B (CTB) or replication-defective HSV-LacZ at E15 (Fig. 2A). Spinal neurons retrogradely-labelled from the cerebellum consist of the double crossed VSCT and the ipsilaterally projecting DSCT neurons. However, dI2 neurons, double labelled by genetic targeting and back labelling from the cerebellum, are all VSCT neurons, since dI2 are commissural neurons. Only the large-diameter neurons were co-labeled, most of them in the ventral aspect of lamina VII (Fig. 2D,E,H; S4B,C). Interestingly, many of CTB^+^ or LacZ^+^ neurons were contacted by dI2 axons (Fig. 2F,G,I; S4D,E), suggesting that small-diameter dI2 neurons innervate pre-cerebellar neurons.

Segmental crossing at the lumbar level and re-crossing back to the ipsilateral side at the cerebellum, is a characteristic of the VSCT projection pattern. To measure the proportion of dI2 in VSCT neurons, we labeled VSCT axons with GFP and dI2 axons with cherry (for experimental design see Supp. Fig. S4G,F). The number of axons expressing the reporters at the contralateral superior cerebellar peduncle was scored. Ten percent of the VSCT axons belonged to dI2 (Fig. S4H). Thus, the large diameter dI2s are 10% of the VSCT neurons, consistent with the anatomical observation that VSCT comprise a heterogeneous population of interneurons (Jankowska and Hammar, 2013, Stecina et al., 2013).

### dI2 neurons receive synaptic input from pre-motoneurons and sensory neurons

To assess the synaptic input to dI2 neurons we investigated their synaptic connectivity with known pre-VSCT neurons: dorsal root ganglion (DRG) neurons, inhibitory and excitatory pre-motoneurons and reticulospinal tract neurons. Two genetically defined pre-motoneurons were examined: V1 – an inhibitory pre-motoneuronal population (Bikoff et al., 2016, Gosgnach et al., 2006) and dI1i excitatory INs. A density plot of dI1 synapses shows dI1i terminals within the motoneuron pools (lamina IX) and at lamina VII (Fig. 3C). Co-labeling of motoneurons and dI1 synapses revealed synaptic contacts of dI1i on motoneurons (Fig. S5A-B). This was further supported by co-labeling dI1 with pre-MNs via hindlimb injection of pseudorabies virus (PRV) (Fig. S5C-D).

To identify neurons presynaptic to dI2 cells we initially assessed the likelihood of connectivity by examining spatial overlap of axonal terminals from the presumed presynaptic neurons and the somata of dI2 neurons, and consequently by detection of synaptic boutons on the somatodendritic membrane of dI2 neurons. We have shown previously that genetic targeting of synaptic reporter in chick spinal cord neurons, yields overlapping expression of the reporter and endogenous pre-synaptic proteins (Hadas et al., 2014). This observation was confirmed using synaptic reporter expressed in dI1 and cytoplasmic reporter expressed in dI2 (Fig. 3C’). The dI2, DRG, V1, and dI1 neurons were labeled using specific enhancers (Fig. S1A). General premotor INs were labeled by injection of PRV-cherry virus into the ipsilateral hindlimb musculature (Hadas et al., 2014) (Fig. 3B).

**Figure 3:**
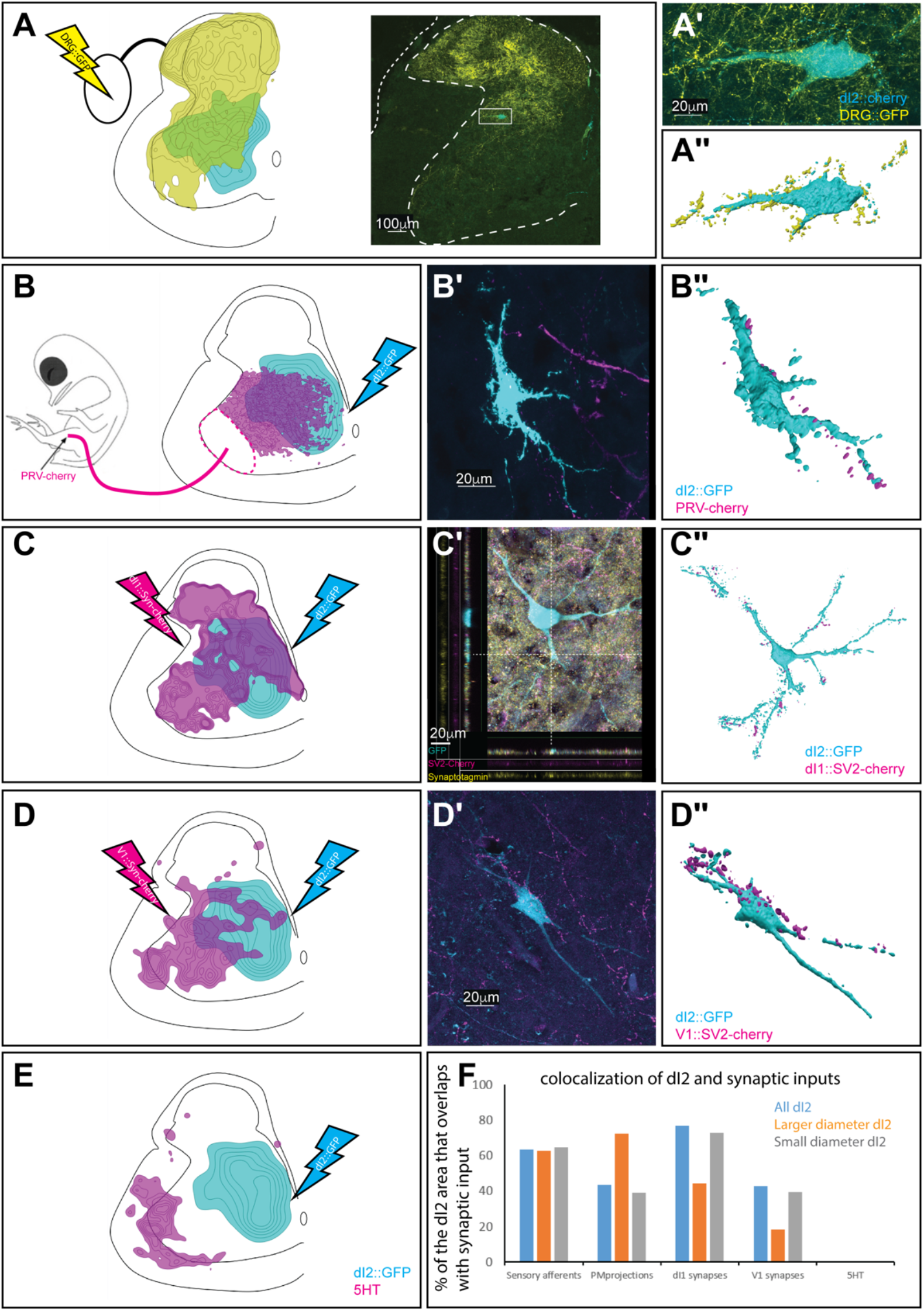
Synaptic inputs to dI2 neurons. Schematic representations of the experimental design for labeling dI2::GFP or dI2::cherry INs (cyan) and potential sources of synaptic inputs (yellow or magenta), supplemented by cell soma density of dI2 INs and the synaptic densities are illustrated in A, B, C, D, E. The density values presented are 10-80%, 20-80%, 25-80%, 30-50% and 20-80%, respectively. Examples of dI2 neurons contacted by axons or synaptic boutons are shown in A’, B’, C’, and D’; and their 3D reconstruction in A’’, B’’, C’’, and D’’. Genetic labeling was attained using specific enhancers (Fig. S1) electroporated at HH18. **A.** DRG neurons form contacts on dI2 neurons. Inset in A: cross section of and E17 embryo at the crural plexus level of the lumbar cord. A dorsally located dI2 neuron contacted by numerous sensory afferents, magnified in A’ and 3D-reconstructed in A’’. (*N*=18 sections) **B.** Premotoneurons form contacts on dI2 neurons. dI2 neurons were labeled at HH18. At E13, PRV virus was injected to the leg musculature, and the embryo was incubated until premotoneuronal infection (39 hours) (*N*=34 sections). **C.** dI1 neurons form synapses on dI2 neurons. (*N*=8568 synapses). C’: A representative SV2::cherry synapse on dI2 dendrite, positive for synaptotagmin. Demonstrated by a horizontal and vertical optical sections in Z-axis (see cursors and color channels). **D.** V1 neurons form synapses on dI2 neurons. (*N*=1923 synapses). **E.** dI2 neurons do not have 5HT synaptic terminals (*N*=1718 synapses). E17 cross sections of dI2::GFP labeled embryos were stained for 5HT. **F.** Quantification of the overlap area of the different input sources and dI2 neuron densities plots. See Figure S5, S6.

A density profile of the axons of DRG neurons (Fig. 3A, S6A-D) and PRV-labeled pre-MNs (Fig. 3B), and a density profile of synaptic boutons from dI1i (Fig. 3C, S6E), V1 (Fig. 3D), and reticulospinal tract neurons (5HT^+^ synapses) (Fig. 3E, S6F) were aligned to the density plots of the dl2 somata. The overlap between the axonal terminals of DRG neurons, PRV-labeled pre-motoneurons, dI1i and V1 boutons, and the somata of dI2 neurons is evident (Fig. 3A-F, Fig. S6G). To further substantiate these putative synaptic connections, we performed double labeling of DRG, PRV-labeled pre-motoneurons, V1, and dI1i, together with dI2 neurons. Hindlimb pre-motor neurons were labeled by injection of PRV into the hindlimb musculature (Hadas et al., 2014); DRG and dI1 terminal and synapses by cell-type specific enhancer (Fig. S1A); V1 synapses by a double conditional on/off reporter plasmids that enables activation of synaptic reporter in V1 and dI2 utilizing the Foxd3 enhancer and deletion of the synaptic reporter from dI2 neurons utilizing the Ngn1 enhancer (Fig. S1A).

Contact between DRG axons and dI2 neurons was mainly apparent in the dorsal dI2 neurons, while the ventral dI2 neurons received little to no input from DRG neurons (Fig. 3A, S6A,B). Synaptic connections, evaluated by boutons on dI2 dendrites and somata, are apparent from the pre-motor neurons V1 and dI1i (Fig. 3C,D). Serotonergic synapses were found to be concentrated on motoneurons and were not observed on dI2 neurons (Fig. 3E, S6F). Double-labeling of 5HT and dI2 neurons did not reveal any synaptic input. The lack of synaptic serotonergic input may be related to the difference in species, or may suggest that other, non-dI2 VSCT neurons residing adjacent to motoneurons, are contacted by the reticulospinal neurons. The analysis of the synaptic inputs supports the concept that dI2 neurons constitute part of the VSCT. They receive input from sensory afferents, inhibitory and excitatory pre-motoneurons and project to the cerebellum.

### dI2 neurons innervate contralateral lumbar and brachial pre-MNs and dI2 neurons

Axon collaterals of dI2 invade the spinal cord gray matter along the entire length of the spinal cord, as revealed by whole mount staining of spinal cords electroporated with alkaline phosphatase reporter (dI2::AP) (Fig. 4A) and membrane-tethered EGFP (Fig. 4B). The region innervated by dI2 collaterals (arrow at Fig. 4B) overlaps with that of the pre-MNs, V0 and V1 (Lai et al., 2016, Griener et al., 2015), as well as with that of the contralateral dI2 neurons (Fig. 1, Fig. 4B). To assess the potential spinal targets of dI2 neurons, we scored the degree of overlap between of dI2 synapses and dI2 somata (Fig. 4C), the ipsilateral pre-MNs (Fig. 4D), and contralateral pre-MNs (Fig. 4E). The alignment revealed a significant overlap of dI2 synapses with ipsi/contralateral pre-MNs and dI2 neurons (Fig. 4G,H). Co-labeling of synaptic dI2 coupled with labeling of the above neuronal population showed dI2 synaptic boutons on pre-MNs and dI2 neurons at the lumbar level (Fig. 4C-H, S7A-C).

**Figure 4:**
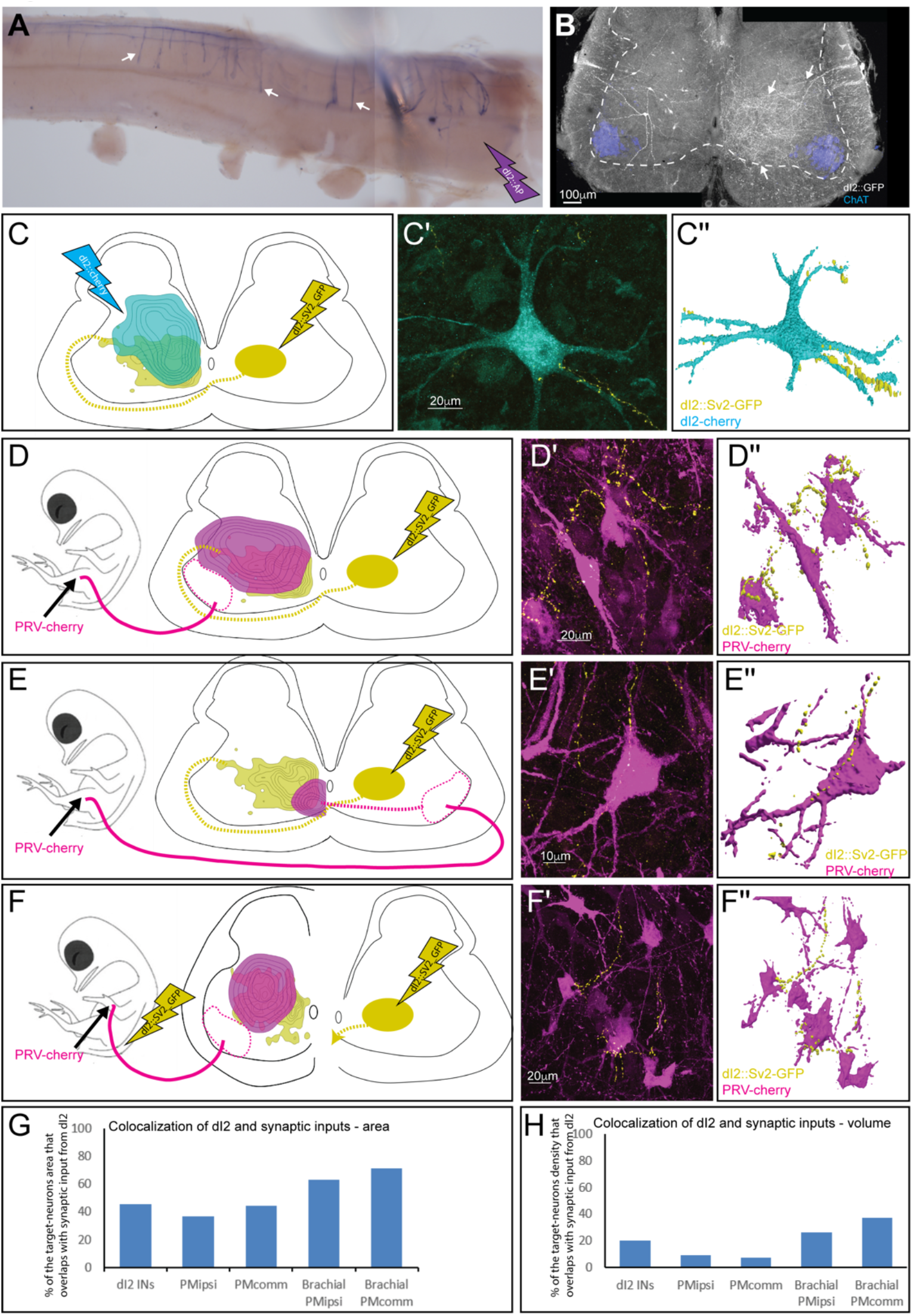
Sp inal synaptic targets of dI2 neurons. **A.** A whole mount staining of spinal cord (thoracic segments) expressing alkaline phosphatase (AP) in dI2 neurons. The lumbar dI2 neurons (not included in the image) were labeled with AP. dI2 axon collaterals project and into the spinal cord (arrows). **B.** Cross section of an E17 embryo at the crural plexus level of the lumbar cord. The axon collaterals (white arrow) penetrating the gray matter of the contralateral side are evident. Schematic representations of the experimental design for labeling synapses (dI2::SV2-GFP, yellow) and potential targets (magenta) supplemented by cell soma density and dI2 synaptic densities are illustrated in C, D, E, and F. Examples of target neurons contacting synaptic boutons of dl2 neurons are shown in C’, D’, E’, and F’; and their 3D reconstruction in C’’, D’’, E’’, and F’’. Genetic labeling was attained using dI2 enhancers (Fig. S1) electroporated at HH18. Pre-MNs were labeled by injection of PRV-cherry into the hindlimbs (D, E) or the forelimb (F) musculature, at E13. The embryos were incubated until pre-MNs infection (39 hours). **C.** dI2 neurons innervate the contralateral dI2 neurons (*N*=4735 synapses and *N*=374 cells, respectively). **D.** dI2 neurons innervate ipsilateral projecting premotoneurons at the sciatic plexus level (*N*=4735 synapses and *N*=936 cells, respectively). **E.** dI2 neurons innervate contralateral projecting premotoneurons at the sciatic plexus level (*N*=4735 synapses and *N*=47 cells, respectively). **F.** dI2 neurons innervate brachial ipsilateral projecting premotoneurons (*N*=2215 synapses and *N*=286 cells, respectively). **G.** Quantification of the overlap area of different synaptic targets and dI2 synapse density plots, as the percentage of overlap between dI2 synapses and the target. **H.** Quantification of the overlap in volume of the different synaptic targets and dI2 synapse density plots, as the percentage of overlap between the synaptic target and dI2 synapses. See Figure S7.

The pattern of dI2 collaterals along the entire rostrocaudal axis (Fig. 4A) suggests that dI2 neurons innervate contralateral pre-MNs and dI2 neurons at multiple levels. To test this, labeling of lumbar dI2 neurons was coupled with labeling of brachial pre-MNs and dI2 soma labeling via wing musculature injection of PRV or electroporation of reporter in brachial dI2 neurons, respectively (Fig. 4F, S7D). dI2-synapses and the putative targets overlapped, and synaptic boutons originated from lumbar level dI2 neurons are apparent on dI2 neurons, and on the contra- and ipsilateral pre-MNs of the wings (Fig. 4F, S7D-F).

The neuronal and synaptic labelling experiments showed that lumbar dI2 neurons innervate the cerebellum, lumbar and brachial pre-MN, and contralateral dI2 neurons (Fig. 6D). Hence, dI2 neurons may relay peripheral and intraspinal information to the cerebellum and to the contralateral lumbar and brachial motor control centers.

### Silencing of dI2 neurons impairs stability of bipedal stepping

The synaptic input to dI2 and their targets, implicate them as relaying information about motor activity to the contralateral spinal cord and the cerebellum. Thus, we hypothesized that manipulation of their neuronal activity may affect the dynamic profile of stepping.

To study the physiological role of dI2 neurons, we silenced their activity using bilateral targeting of the tetanus toxin (*TeTX*) gene, to lumbar dI2s. EGFP was co-targeted in a 2/1 TeTX/EGFP ratio. EGFP expression in dI2 neurons and non-electroporated chicks was used as a control. In order to maximize the number of targeted dI2 neurons, we combined genetic targeting with the *Foxd3* enhancer and spatial placement of the electrodes at the dorsal lumbar spinal cord (Fig. S1A). Embryos were electroporated at E3. Upon hatching, chicks were trained for targeted over-ground locomotion.

To test whether silencing of dI2 impairs post-hatching development and muscle strength, at P8 the weight of the chicks was measured and the foot grip power evaluated (see below). Gait parameters of 4 controls and 5 *TeTX* treated chicks were measured while chicks were walking toward their imprinting trainer along a horizontal track (6-20 walking sessions, 5-8 strides each, per chick). Following the experiments, chicks were scarified and their spinal cords were removed and processed for immune-detection of the efficacy of the electroporation (Supplementary table S1).

The weight of all chicks was comparable within the range of 144.7±12.1 gr (Supp. Table S1). As a functional measure of foot grip, we tested the ability of the chicks to maintain balance on a tilted meshed surface. *TeTX*-manipulated chicks and control chicks, maintained balance on the titled surface up to 63–70^0^, with no apparent statistically significant differences (Supplementary Table S1, Supplementary statistics). Thus, manipulation of dI2 neuronal activity did not impair post-hatching development and muscle strength.

Analysis of over ground locomotion of the control and *TeTX* treated chicks revealed no significant differences is swing velocity and striding pattern. An 180^0^ out-of-phase pattern was found during stepping in all the manipulated and the control chicks (Fig. S8A, table 1, Supplementary statistics). However, substantial differences were scored in stability parameters: *TeTX* chicks exhibited whole-body collapses during stepping (Fig. 5B,C, Fig. 6A), wide-base stepping (table 2), and variable limb movements (Fig.5 D,E, Fig. 6B,C, Fig. S8B,C, tables 3, Supplementary statistics).

**Figure 5:**
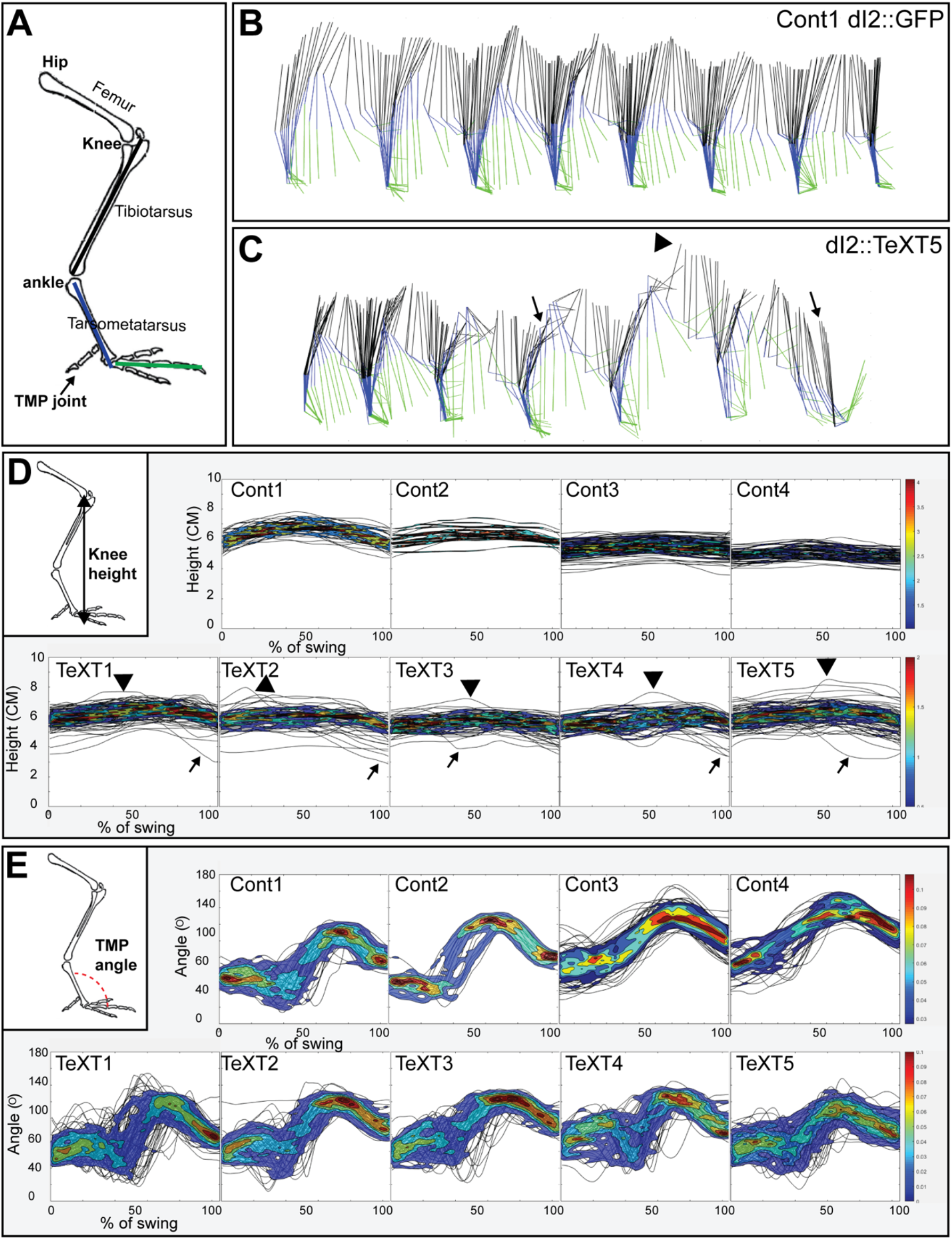
Kinematic analysis of locomotion in post-hatching chicks following neuronal silencing of dI2. **A.** Schematic illustration of chick hindlimb joints (bold) and bones (regular). The knee joint connects between the femur and the tibiotarsus, the ankle connects the tibiotarsus and the tarsometatarsus which connects to the phalanges by the tarsometatarso-phalangeal joint (TMP). During the swing phase of birds, the ankle flexion leads to foot elevation, while the knee is relatively stable. **B, C.** Stick diagrams of stepping in a control chicken d2::GFP (B) and in a d2::TeTX chicken (C). Arrows indicate falls and overshoots are denoted by arrowheads. **D.** Overlays of knee height (demonstrated in insert) trajectories during the swing phase in all analyzed steps of each of the control and *TeTX* treated P8 hatchlings are shown superimposed with the respective 20%–80% color coded density plots as a function of the percentage of swing (see Text and Materials and Methods). Arrows indicate falls and overshoots are indicated by arrowheads. **E.** Overlay of angular trajectories of the TMP joint (demonstrated in insert) during the swing phase in all analyzed strides of each of the control and *TeTX* treated P8 hatchlings are shown superimposed with the respective 20%–80% color coded density plots as a function of the percentage of swing (see Text and Materials and Methods).

**Figure 6:**
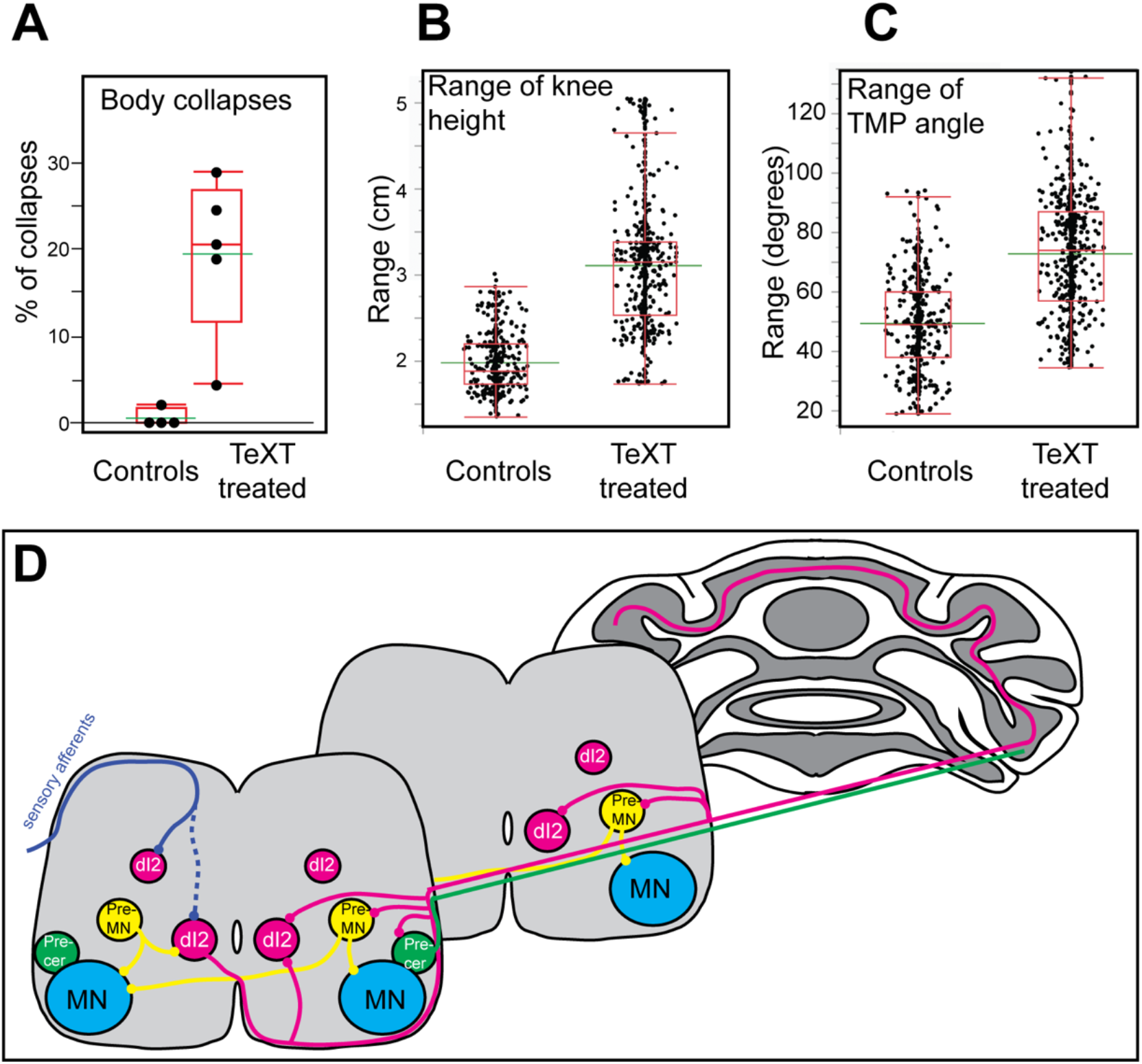
Increased inter-stride variability in TMP angle and knee height in *TeTX*-treated chicks. **A.** The percentage of steps with body collapses in the controls and *TeTX* manipulated hatchlings (n=4 and n=5, respectively). p-value<0.0001 (Z-test). See table 3 for single chick fall proportions. **B.** Analysis of the mean range of knee height changes during the swing phase of control and TeTX treated chicks (n=4 and n=5, respectively). p-value<0.0001 using a T-test allowing different variances. See Figure S8B and table 3 for single chick data and statistical analysis details. **C.** Analysis of the mean range of TMP angular excursions during the swing phase of control- and TeTX-treated chicks. (n=4 and n=5, respectively). p-value<0.0001 Watson and Williams F test. See Figure S8C and table 3 for single chick data and statistical analysis details. **D.** Schematic illustration showing the connectome of lumbar dI2 neurons. dI2s (magenta) receive synaptic input from sensory afferents (solid blue line indicates massive synaptic input and dashed blue line indicates sparse innervation), Inhibitory and excitatory premotoneurons (yellow), and from the contralateral lumbar dI2. dI2s innervate the contralateral lumbar and brachial premotoneurons (both commissural and ipsilateral projecting premotoneurons are innervated by dI2), the lumbar and brachial contralateral dI2, lumbar pre-cerebellar neurons (green) and the granule neurons in the cerebellum.

**Table 1:**
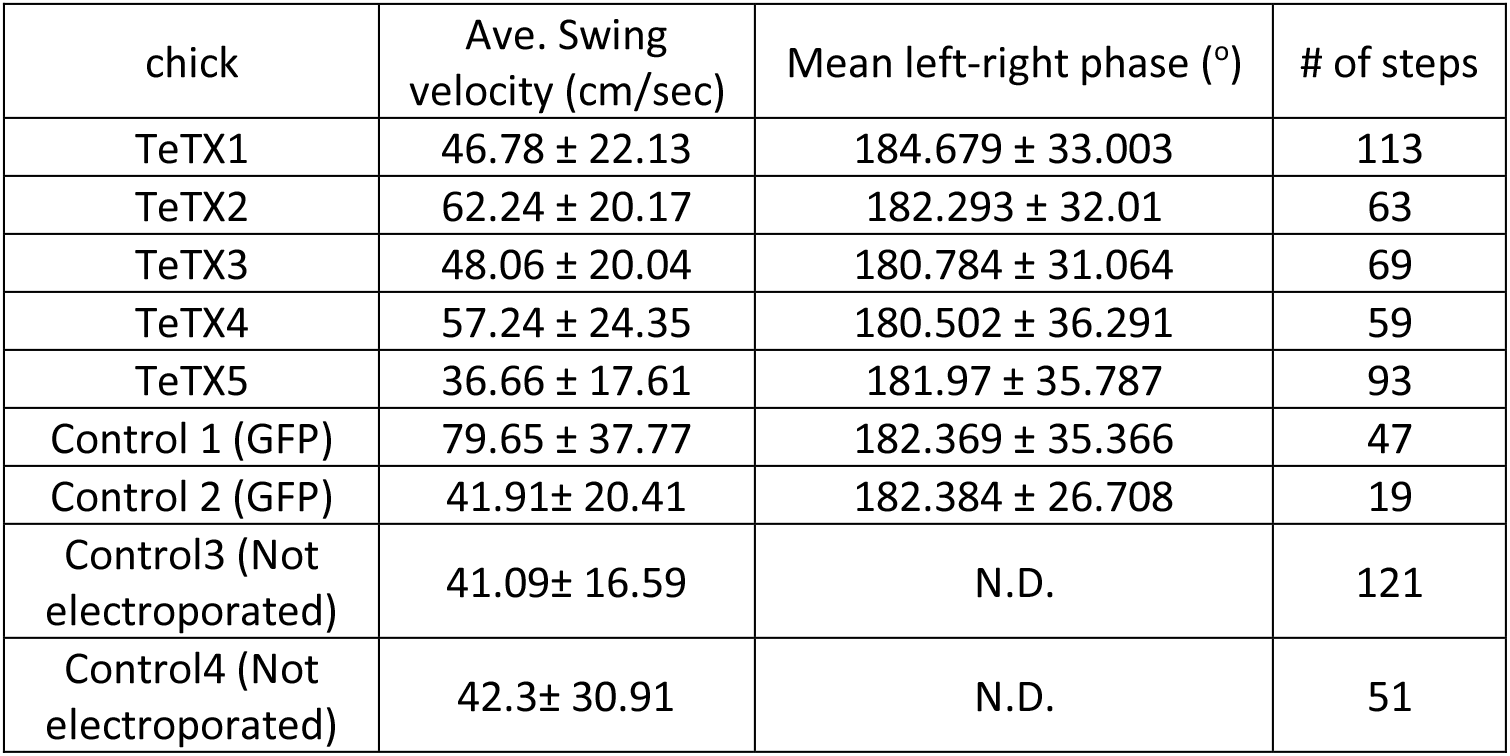
Stride velocity and left-right phase in control and TeTx manipulated chicks Swing velocity and left-right phase were measured and calculated as described in the Materials and Methods. The Watson and Williams F test of the phase data (circular ANOVA) were not statistically significant (See also Supplementary statistics).

| chick | Ave. Swing velocity (cm/sec) | Mean left-right phase (°) | # of steps |
| --- | --- | --- | --- |
| TeTX1 | 46.78 ± 22.13 | 184.679 ± 33.003 | 113 |
| TeTX2 | 62.24 ± 20.17 | 182.293 ± 32.01 | 63 |
| TeTX3 | 48.06 ± 20.04 | 180.784 ± 31.064 | 69 |
| TeTX4 | 57.24 ± 24.35 | 180.502 ± 36.291 | 59 |
| TeTX5 | 36.66 ± 17.61 | 181.97 ± 35.787 | 93 |
| Control 1 (GFP) | 79.65 ± 37.77 | 182.369 ± 35.366 | 47 |
| Control 2 (GFP) | 41.91± 20.41 | 182.384 ± 26.708 | 19 |
| Control3 (Not electroporated) | 41.09± 16.59 | N.D. | 121 |
| Control4 (Not electroporated) | 42.3± 30.91 | N.D. | 51 |

**Table 2:** Maximum stride width in control and TeTx manipulated chicks Stride width was measured as described in Materials and Methods. One-way ANOVA followed by Dunnett’s.

| chick | maximum stride width (cm) | # of steps |
| --- | --- | --- |
| TeTX1 | 5.11 ± 1.89 | 97 |
| TeTX2 | 5.32 ± 1.38 | 36 |
| TeTX3 | 4.5 ± 1.01 | 27 |
| TeTX4 | 4.9 ± 1.16 | 49 |
| TeTX5 | 5.82 ± 1.71 | 110 |
| Control 8 | 4.15 ± 1.07 | 137 |
| Control 9 | 4.32 ± 1.32 | 115 |

**Table 3:**
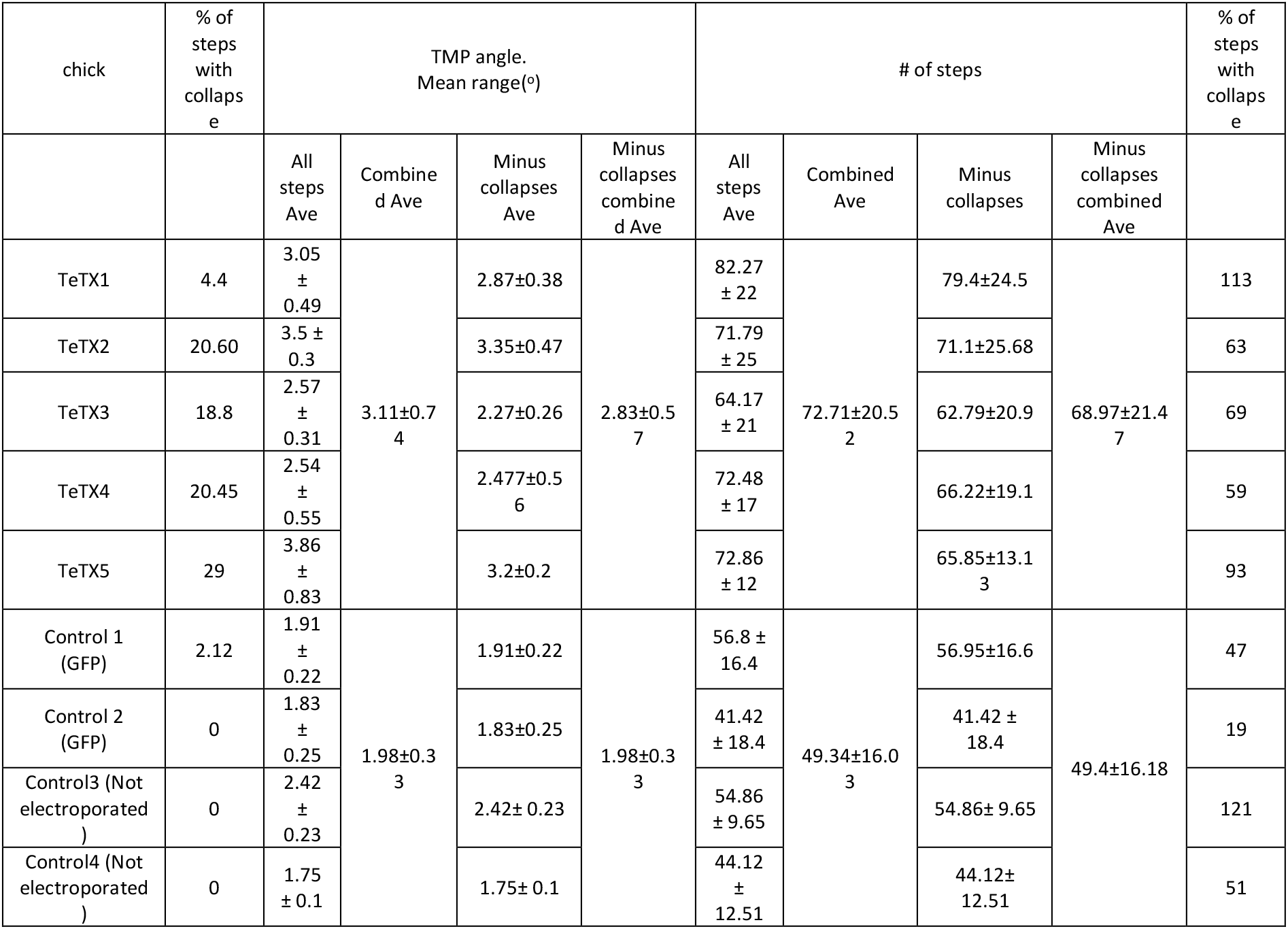
Collapses, knee height and TMP angle ranges in control and TeTx manipulated chicks. Analysis of the range between the highest and lowest point of knee height and the highest and lowest angle of the TMP joint in all steps before and after subtraction of collapsed steps. When combined – the difference is statically significant (p-value<0.0001 for both metrics, using a T-test allowing different variances).

#### Whole-body collapses

A collapse was scored as a decline of the knee height below 85% of the average knee height at the stance phase of the step (arrow in Fig. 5C). Collapses were usually followed by over-extensions (arrow head in Fig. 5C). We measured the number of collapses in 50–190 steps. In control chicks, collapses occurred in 0.53±0.92% of the steps. In *TeTX* manipulated chicks we observed collapses in 19.46±8.3% of the steps, significantly different from the controls (Fig. 6A, Supplementary statistics). The collapses and over-extensions were also manifested in the profiles of the knee height trajectory during the swing phase (Fig. 5D).

#### Wide base stepping

Wide-base stance is typical of an unbalanced ataxic gait. The stride width was measured between the two feet during the double stance phase of stepping. The mean stride in *TeTX*-manipulated chicks 1,2,4, and 5 was significantly wider than the control chicks, while the width in *TeXT3* was similar to the controls (Table 2, Supplementary statistics).

#### Variable limb movements

In stable gait, limb trajectories are consistent from stride to stride. For point by point comparisons of the trajectories in knee height and angles of the TMP joint during the swing phase of stepping in control and TeTx manipulated chicks, the swing phase was normalized using 126 consecutive epochs, and the data were displayed as a function of the percentage of swing. Plots of the knee height and TMP angle trajectories during the normalized swing in all the analyzed steps of each chick are shown superimposed in Fig. 5D and Fig. 5E, respectively. These data demonstrate that the range of changes in *TeTX*-manipulated chicks was higher comparing to control chicks.

Further analyses revealed that overall the control group shows lower Knee-height and TMP angle ranges than the TeXT-treated group, even though there are differences within groups (Fig. S8B,C). The average knee height range of the combined control chicks (1.981±0.33) is significantly lower than the range of the combined TeXT treated chicks (3.109±0.74) (Fig. 6B, Supplementary statistics). Similar comparison of the combined ranges of angular excursions of the TMP joint during the normalized swing revealed that the average angle of the controls group (49.34±16.03) is significantly lower than the average of the TeXT-treated chicks (77±22.149). (Fig. 6C, Supplementary statistics).

Since the increased range of changes could be due to the effects of the substantial increase in body collapses during stepping (Fig. 6A, see also Fig. 5), we excluded steps featuring whole body collapses and reanalyzed the data. The data summarized in table 3, shows that the significant difference between controls and the *TeXT*-treated chicks in the range of the knee height and the TMP angle excursions was maintained. Thus, the increase in irregularity in the *TeXT*-treated chicks is not only due to the body collapses of the *TeXT*-treated chicks. Overall, the kinematic parameters of the *dI2-TeXT-treated* chicks demonstrate a reduction in stability during locomotion, featuring a possible role of dI2 in stabilizing bipedal stepping.

## Discussion

The VSCT is thought to provide peripheral and intrinsic spinal information to the cerebellum in order to shape and update the output of spinal networks that execute motor behavior. The lack of genetic accessibility to VSCT neurons hampers elucidation of their role in locomotion. Using genetic toolbox to dissect the circuitry and manipulate neuronal activity in the chick spinal cord, we studied spinal interneurons with VSCT characteristics. The main findings in our study are that dI2 neurons in the chick lumbar spinal cord are commissural neurons that innervate premotor neurons at the contralateral lumbar and brachial spinal levels and granule neurons in the ipsilateral cerebellum. Hence, subpopulation of dI2 form a part of the avian VSCT. Targeted silencing of dI2 neurons leads to impaired stepping in P8 hatchlings. We described the spatial distribution of sub-populations of dI2 neurons, deciphered their connectomes, mapped the trajectory of their projection to the cerebellum, and suggested possible mechanisms for the perturbed gait resulting from their genetic silencing, as discussed below.

Using the intersection between genetic drivers and spatially restricted delivery of reporters to define lumbar and brachial neurons, we have identified several targets of dI2 lumbar neurons. Ipsilateral lumbar dI2 neurons innervate contralateral lumbar dI2 neurons as well as commissural and non-commissural lumbar pre-motoneurons. This connectivity may affect the bilateral spinal output circuitry at the lumbar cord (e.g. (Bras et al., 1988, Jankowska and Hammar, 2013)). Moreover, the ascending axons of lumbar dI2s, give off grey matter collaterals innervating contralateral dI2s and commissural and non-commissural pre-motoneurons throughout the brachial spinal cord (Fig. 6D). Therefore, lumbar dI2 neurons may also contribute to the inter-enlargement coupling described between the limb and wing moving segments of the spinal cord (e.g. (Valenzuela et al., 1990, Ruder et al., 2016) for forelimb to hindlimb coupling connectivity in mice).

We demonstrated that lumbar dI2s receive sensory innervation, pre-motor inhibitory and excitatory innervation, and innervation from contralateral lumbar dI2s (Fig. 6D). Thus, lumbar dI2 neurons can provide the cerebellum and contralateral premotor neurons with proprioceptive information, copies of motor commands delivered from the ipsilateral pre-motoneurons, and integrated information from contralateral dI2 neurons (Fig. 6D).

The unclear genetic origin of physiologically equivalent lumbar VSCT neurons, prevented better understanding of their role in hindlimb locomotion. Our wiring and neuronal-silencing studies implicated dI2 as a significant contributor to the regularity and stability of locomotion in P8 hatchlings. The kinematic analysis of TeTX treated hatchlings, revealed imbalanced locomotion with occasional collapses, increased stride variability, wide base stepping, and variable limb movements during stepping.

The mechanisms accounting for the impaired stepping following dI2 silencing are still unknown. One of the possible mechanisms is that silencing the dI2s perturbs the delivery of peripheral and intrinsic feedback to the cerebellum, leading to unreliable updating of the motor output produced by the locomotor networks, thereby impairing the bipedal stepping. Another possible mechanism is based on the similarity of the gait instabilities of TeTX treated hatchlings to ataxic motor disorders. Mammalian VSCT neurons receive descending input from reticulospinal, rubrospinal and vestibulospinal pathways (Bras et al., 1988, Jankowska and Hammar, 2013). Neurons from the lateral vestibular nucleus, have been reported to innervate extensor motoneurons at the lumbar level, as well as interneurons residing at the medial lamina VII (Murray et al., 2018), exactly at the location where dI2 neurons were found to reside in our study. Thus, the vestibulospinal tract may convey input directly to the ipsilateral motoneurons, and indirectly to contralateral motoneurons through dI2 neurons that innervate contralateral pre-motoneurons. This way silencing of dI2 neurons is expected to interfere with the descending regulation of the stability of the bipedal gait.

The local spinal connections between dI2 and the contralateral premotor neurons and contralateral dI2, may serve an important component of coordinated limb movements. dI2 synapses were found on both ipsilateral and contralaterally projecting pre-motorneurons, in their segmental level and in the brachial level, which regulates the movement of the wings. Thus, dI2 neurons may affect the motor output of the contralateral side of the cord, and also the ipsilateral side by contacting commissural pre-motorneurons.

In summary, our mapping studies of dI2 neurons and their connectomes followed by characterization of the effects of their silencing on bipedal stepping, offer new insights on the function of dI2 neurons in vertebrates. We suggest that lumbar dI2 neurons are not merely used to relay sensory and intrinsic spinal networks information to the cerebellum, but also act as active mediators of motor functions at the lumbar segments and at the wing controlling brachial segments of the spinal cord. Further circuit deciphering studies of the constituents of sub-populations of dI2s, their targets, and their descending inputs are required to extend our understanding of the function of dI2 subpopulations in motor control of movements.

## Materials and Methods

### Animals

Fertilized White Leghorn chicken eggs (Gil-Guy Farm, Israel) were incubated in standard conditions at 38 °C. All experiments involved with animals were conducted in accordance with the designated Experiments in Animals Ethic Committee policies and under its approval.

### 3D reconstruction and density plots analysis

The codes for both 3D reconstruction and the density plots analysis were written in Matlab. The density plots were generated based on cross section images transformed to a standard form. The background was subtracted, and the cells were filtered automatically based on their soma area or using a manual approach. Subsequently, two-dimensional kernel density estimation was obtained using the MATLAB function *“kde2d”*. Finally, unless indicated otherwise, a contour plot was drawn for density values between 20% and 80% of the estimated density range, in six contour lines.

### *In-ovo* electroporation

A DNA solution of 5 mg/mL was injected into the lumen of the neural tube at HH stage 17–18 (E2.75-E3). Electroporation was performed using 3 × 50 ms pulses at 25–30 V, applied across the embryo using a 0.5-mm tungsten wire and a BTX electroporator (ECM 830). Following electroporation, 150–300 μL of antibiotic solution, containing 100 unit/mL penicillin in Hank’s Balanced Salt Solution (Biological Industry, Beit-Haemek) was added on top of the embryos. Embryos were incubated for 3–19 days prior to further treatment or analysis.

### Immunohistochemistry and *In situ* hybridization

Embryos were fixed overnight at 4°C in 4% paraformaldehyde/0.1 M phosphate buffer, washed twice with phosphate buffered saline (PBS), incubated in 30% sucrose/PBS for 24 h, and embedded in OCT (Scigen, Grandad, USA). Cryostat sections (20 μm). Sections were collected on Superfrost Plus slides and kept at −20°C. For 100-μm sections, spinal cords were isolated from the fixed embryos and subsequently embedded in warm 5% agar (in PBS), and 100μm sections (E12–E17) were cut with a Vibratome. Sections were collected in wells (free-floating technique) and processed for immunolabeling.

The following primary antibodies were used—rabbit polyclonal GFP antibody 1:1000 (Molecular Probes, Eugene, Oregon, USA), mouse anti-GFP 1:100, Goat anti-GFP 1:300 (ABcam), rabbit anti-RFP 1:1000 (Acris), goat ChAT antibody 1:300 (Cemicon, Temecula, CA, USA), mouse anti-synaptotagmin antibody 1:100 (ASV30), mouse anti-Lhx1/5 1:100 (4F2), mouse anti-FoxP4 1:50 (hybridoma bank, University of Iowa, Iowa City, USA), mouse anti Brn3a 1:50 (Mercury), rabbit anti-Pax2 antibody 1:50 (ABcam), chicken anti-lacZ antibody 1:300 (ABcam), rabbit anti Clabindin 1:200 (Swant), rabbit VGUT2 antibody (Synaptic Systems, Göttingen, Germany), goat anti FoxP2 1:1000 (ABcam), goat anti FoxP1 1:100 (R&D Systems) and rabbit anti 5HT 1:100 (Immunostar). The following secondary antibodies were used: Alexa Fluor 488/647-AffiniPure Donkey Anti mouse, rabbit and goat (Jackson) and Rhodamin Red-X Donkey Anti mouse and rabbit (Jackson). Images were taken under a microscope (Eclipse Ni; Nikon) with a digital camera (Zyla sCMOS; Andor) or a confocal microscope (FV1000; Olympus).

*In situ* hybridization was performed as described (Avraham O et al., 2010). The following probes were employed: Foxd3, vGlut2 and GAD1 probes were amplified from a cDNA of E6 chick embryo using the following primers. Foxd3: forward-TCATCACCATGGCCATCCTG and Reverse -GCTGGGCTCGGATTTCACGAT. vGlut2: forward -GGAAGATGGGAAGCCCATGG and Reverse -GAAGTCGGCAATTTGTCCCC. GAD1: forward-TCTCACCTGGAGGAGCCATC and Reverse -CCTGAGGCTGATATCCAACC. T7 RNApol cis-binding sequence was added to the reverse primers.

### AP staining

The treated embryos were fixed with 4% paraformaldehyde–PBS for 24 h at 4°C, and washed twice with PBS for 30 min at 4°C. The fixed embryos were incubated at 65°C in PBS for 8 to 16 h to inactivate the endogenous AP activity. The treated embryos were washed with 100 mM Tris–Cl (pH 9.5) containing 100 mM NaCl and 50 mM MgCl2, and the residual Placental alkaline phosphatase activity was visualized by incubating the embryos with NBT/BCIP (Roche) in the same buffer at 4°C for 24 h. After extensively washing the embryos with PBS–5 mM EDTA, the spinal cord was imaged.

### Analysis of Left-right phase

Stride duration was measured as the time from ‘Right toe-off’/foot-off’ to the next Right ‘toe-off’ (as a complete stride cycle for the right leg), and the ‘half-cycle’ duration as the time of Right-toe off to the time of Left toe off. the following formula was used to calculate the phase: ((LeftToeOff_1 – RightToeOff_1)/ (RightToeOff_2 - RightToeOff_1))*360.

### PRV infection

We used two isogenic recombinants of an attenuated PRV strain Bartha (PRV Bartha) that express enhanced GFP (PRV152) and monomeric red fluorescent protein (PRV614). The viruses were harvested from Vero cell cultures at titers 4 × 108, 7 × 108 and 1 × 109 plaque forming units (pfu/mL), respectively. Viral stocks were stored at −80°C. Injections of 3 μL of PRV152 or PRV614 were made into the thigh musculature of E13 or E14 chick embryos, using Hamilton syringe (Hamilton; Reno, NV, USA) equipped with a 33-gauge needle. The embryos were incubated for 36–40 h and sacrificed for analysis. For spinocerebellar projecting neurons labeling, we used a replication defective HSV (TK^−^) that contains a lacZ reporter. The virus was injected *in ovo* into the cerebellum of E12-15 embryos, and the embryos were further incubated for 40–48. Alternatively, cholera toxin subunit B (CTB) conjugated to Alexa Fluor™ 647 (ThermoFisher) was used for the same purpose, and was injected to the cerebellum of E12–15 embryos together with the virus for visualization of both cerebellar projecting neurons and upstream neurons.

### Force test

The muscle strength was evaluated using the measurement of the angle of the fall from a ladder with a gradually increasing-angle. This test was repeated for each chicken at least 3 times, and the average falling angle was calculated.

### Behavioral tests and analysis

The embryos were bilaterally electroporated, and were then allowed let to develop and hatch in a properly humidified and heated incubator. Afterwards, within 32 hours post hatching, the hatchling chicks were imprinted on the trainer. The P8 chicks were filmed in slow motion (240 fps) freely walking (side and top views). The following parameters were scored: 1) weight, 2) foot grip power, 3) kinematics parameters during overground locomotion: a) swing velocity, b) swing and stance duration, c) phase of footfalls, d) height of knee and tarsometatarso-phalangeal (TMP) joints, e) angles of the TMP and ankle joints, and f) stride width (distance between feet during the double stance phase).

Using a semi-automated Matlab-based tracking software (Hedrick, 2008), several points of interest were encoded. The leg joints as well as the eye and the tail were tracked. The position of these reference points was used for computational analysis using an in-house Matlab code for calculating different basic locomotion parameters (e.g. stick diagrams, velocity, joints trajectory, angles, range, and elevation), steps pattern, and degree of similarity (Haimson B et al. *in preparation*). Dunnett’s test (Dunnett, 1955) was used to perform multiple comparisons of group means following One-way ANOVA. Circular statistics was used for analyses of angular data. Circular statistics was used for analyses of angular data, utilizing Oriana (KCS, version 4).

## Supporting information

Haimson et sl., Supplementary Data

## Acknowledgements

The authors thank Haya Falk for PRV purification; Alona Katzir, Cole Bendor, Mevaseret Avital, Sapir Shevah, Eitan Yisraeli, Ruth Segal, Fedaa Bazan and Eden Kimchi for technical assistance; Michael O’Donovan for comments on the manuscript. This work was supported by grants to AK from the Israel Science Foundation (grant No. 1400/16), The US-Israel Binational Science Foundation (grant No. 2017/172) and the Avraham and Ida Baruch endowment fund.

## Author contributions

B.H., A.Klar and ALT designed research; B.H., and Y.H. preformed research; B.H., and M.D. generated the kinematic analysis tools; Y.C. assisted in the post hatching analysis; A. Kania provided reagents and comments on the research; B.H., A.Klar and ALT analyzed the data; B.H., A.Klar and ALT wrote the paper.

