## Supplementary material for "Spinal dI2 interneurons regulate the stability of bipedal stepping": Haimson et sl., Supplementary Data

Supplementary information

**Supplementary Figure legends**

Figure S1: Targeting, reporters, and activity modifiers, used in the study.

**A.** The different DNA constructs used in this study. For targeting expression in dI1 the EdI1 enhancer was used. For targeting dI2, intersection between *Ngn1* and *Foxd3* enhancers was attained. An Isl1 enhancer was used for targeting expression in DRG neuron (Uemura et al., 2005). **B.** A table summarizing the axonal projection, neurotransmitter identity, soma location, and the existence of synaptic connections between pre-MNs and spinal interneuron populations. Based on previous studies (Jessell, 2000, Alaynick et al., 2011, Lai et al., 2016).

Figure S2: Differential expression of transcription factors in dI2 neurons.

**A.** Pre-migratory dI2 neurons are $Lhx1^{+}Pax2^{-}$ (arrows). Cross section of the chick E5 neural tube, expressing GFP in dI2 neurons and immunostained for Lhx1 (yellow arrows) and Pax2.

**B.** Pre-migratory dI2 neurons express *FoxD3* (red arrow), and downregulate its expression upon ventral migration (yellow arrows). Cross section of a chick neural tube at E5, double labeled for dI2 dI2::GFP) and processed for *in situ* hybridization for *FoxD3* probe.

**C.** Genetic heterogeneity in dI2 neuron population. Only a small subset of dI2 neurons expresses the genes *FoxP2* and *FoxP4*. The arrow indicates a $FoxP2^{+}/FoxP4^{+}$ dI2::GFP neuron. Cross section of chick E6 neural tube expressing GFP in dI2 INs and immunostained for FoxP2 and FoxP4.

**D**. A small fraction of dI2 neurons express Pax2, an indicator of inhibitory phenotype. Cross section of chick E6 neural tube expressing GFP in dI2 INs and immunostained for Lhx1 and Pax2. The arrow points to a dI2::GFP/$Lhx1^{+}/Pax2^{+}$ neuron.

Fig. S3: dI2s neurotransmitter phenotype and soma localization.

**A-D.** Most of dI2 neurons are excitatory. dI2 neurons expressing GFP together with *in situ* hybridization using the *Vglut2* probe (A) or the *GAD1* probe (C), and the corresponding quantification of the double-labeled neurons (B and D, respectively).

**E.** Density plot of dI2 soma in the crural plexus segments (*N*=551 cells) (left). Density plot of $dI2_{large}$ (magenta) and $dI2_{small}$ (yellow) neurons in the crural plexus segments (*N*=48 and *N*=502 cells, respectively. N=2 embryos) (right).

**F.** Density plot of dI2 soma in the brachial segments (*N*=66 cells, N=2 embryos)

Figure S4: dI2s densities at the crural plexus level, innervation of deep cerebellar nuclei and projection to the cerebellum.

**A.** dI2 synapses in the deep cerebellar nuclei. A cross section of chick E17 cerebellum. dI2 synapses (magenta), synaptotagmin (cyan). **B.** Density plot of dI2 and precerebellar neurons at the crural plexus segments (*N*=551 and *N*=652 cells, respectively, N=2 embryos). **C.** Quantification of the overlap in area and volume of the two density plots. **D.** Density plot of dI2 synapses and precerebellar neurons at the crural plexus segments (*N*=2543 synapses with density values 25-80% and *N*=652 cells with density values 10-90%, respectively). **E.** Quantification of the overlap in area and volume of the two density plots. **F-G.** dI2 proportion from all VSCT neurons. GFP expression was activated by a removal of a single STOP cassette via Cre-recombinase, and cherry expression was activated by removal of two STOP cassettes via Cre and FLPo recombinases. In control experiment (F) the expression of Cre and FLPo is driven by the ubiquitously expressed CAG enhancer/promoter, while expression in dI2 (G) is controlled by CAG::Cre and dI2 enhancer driving FLPo. F'-G': A section at the superior cerebellar peduncle. The ratio between the number of $Cherry^+$ (VSCT or dI2, in F',F'' and G',G'' respectively) and $GFP^+$ (VSCT) axons was calculated. **H.**

Quantification of Cherry$^+$/GFP$^+$ in each experiment. In control electroporation, the ratio between cherry and GFP is 0.52±0.059 (blue box). Cherry expressing dI2 axons are 0.05±0.01 (Orange box). Thus, considering the 50% efficiency of double versus single conditional removal of STOP signal, the amount of dI2 axons in 9.61%.

(F and G, N=2 and 3 embryos, respectively).

Figure S5: dI1i are pre-motoneurons

**A, B.** Schematic illustration of the strategy for studying the potential innervation of motor neurons by dI1 neurons (A). dI1 neurons were targeted by dI1 specific enhancer (Fig. S1A). At E15 synaptic boutons, labeled with SV2-GFP, were studied on cross sections. Synaptic reporter (cyan) was expressed in dI1 neurons. dI1i boutons contacting Chat$^+$ motorneurons (magenta) are apparent (B).

**C, D.** Schematic representation of the experimental design for co-labeling dI1 and pre-motoneurons, supplemented by cell soma densities of dI1 neurons (cyan, *N*=643 cells) and pre-motoneurons (magenta, N=936 cells). dI1 neurons were labeled at HH18. At E13, PRV virus was injected to the leg musculature, and the embryo was incubated until infection of pre-motoneurons (35 hours) (C). In cross section of E13, PRV-cherry (magenta) is detected in motor neurons (Chat$^+$, yellow) and pre-motoneurons (D). An example of dI1 neurons co-expressing GFP and PRV-cherry shown in D'.

Figure S6: Input of 5HT and dI1 to dI2 neurons at the level of the crural plexus

**A, B.** Sparse sensory innervation of ventrally located dI2 neurons. Cross section of an E17 embryo at the lumbar spinal cord (crural plexus level). A ventrally located dI2 neuron is sparsely contacted by sensory afferents (A), magnified in B. **C, D.** Density plots of $dI2_{large}$ (magenta, *N*=48) and $dI2_{small}$ (yellow, *N*=502) neurons and sensory afferents (*N*=18 sections, with density values 10-80%) in the crural plexus segments. **E.** dI1 neurons form synapses on dI2 neurons (*N*=13369 synapses with density values 50-90%). Example of dI1 boutons (magenta) on a dI2 neuron (cyan) (E') and its 3D reconstruction in (E'').

**F.** dI2 are not contacted by 5HT synaptic terminals (*N*=2754 synapses with density values 25-85%). E17 cross sections of dI2::GFP labeled embryos were stained for 5HT.

**G.** Quantification of the overlap in volume of the different input sources and dI2 neurons densities plots.

<u>Figure S7</u>: Spinal targets of dI2

Schematic representations of the experimental design for labelling synapses (dI2::SV2-GFP, yellow) and potential targets (magenta or cyan), supplemented by cell soma density. The targets and the dI2's synaptic densities are illustrated in A, B, C and E.  Examples of target neurons contacted by dI2's synaptic boutons are presented in A', B', C' and E'; and their 3D reconstruction in A'', B', C'' and E''. Genetic labelling was attained using dI2 enhancers (Supp Fig. S1) electroporated at HH18. Pre-MNs were labelled by injection of PRV-cherry into the hindlimbs (D, E) or the forelimb (F) musculature, at E13. Embryo was incubated until pre-motorneurons infection (39 hours).

**A.** dI2 innervate ipsilateral projecting pre-motoneurons at the crural plexus level (*N*=2543 synapses and *N*=250 cells, respectively).

**B.** dI2 innervate contralateral projecting pre-motoneurons at the crural plexus level (*N*=2543 synapses and *N*=117 cells, respectively).

**C.** dI2 innervate the contralateral dI2 neurons at the crural plexus level (*N*=2543 synapses and *N*=551 cells, respectively).

**D.** dI2 innervate brachial contralateral projecting pre-MNs (*N*=2215 synapses and *N*=90 cells, respectively).

**E.** dI2 innervate brachial dI2 neurons (N=2215 synapses and *N*=66 cells, respectively).

**F.** Quantification of the overlap in area and volume of the different synaptic targets and dI2 synapse density plots, as percentage of overlap of dI2 synapses with the target.

<u>Figure S8</u>: Locomotion characteristics of control and *TeTX*-treated chicks.

**A.** The mean left-right phase of the control and *TeTX*-treated chicks. The mean phase values are pointed at by the r-vectors (arrows). The Rayleigh critical values (P=0.05) are indicated by blue circles.

**B-C**. The range of knee height and TMP angle (B and C, respectively, see text for details) for each chicken (four controls and five TeTX-treated chicks).

Purple lines –averages of each group, controls and TeXT-treated. Blue lines – average of individual chick. Red lines and boxes – box plot. Black lines – the range of each chick.

In general, a higher inter-stride variability in the *TeTX*-treated chicks is evident.

Supplementary table S1

Weight, force and number of electroporated cells. See statistical tests in Supplementary Statistical analysis tables.

.

| chick | # of dI2::TeTX cells | % of large diameter dI2::TeTX cells | Weight in gr. | Force test angle of fall | |
|---|---|---|---|---|---|
| | | | | Mean ± circ*SD* | *N* |
| TeTX1 | 602 | 5.64 | 139 | 68.41 ± 2.8 | 3 |
| TeTX2 | 124 | 6.45 | 139 | 63.34 ± 1.85 | 3 |
| TeTX3 | 81 | 7.4 | 159 | 65.45 ± 3.35 | 5 |
| TeTX4 | 755 | 7.01 | 148 | 69.29 ± 3.48 | 5 |
| TeTX5 | 769 | 8.71 | 139 | 66.44 ± 3.75 | 3 |
| Control 3 | | | 158 | 64.21 ± 2 | 16 |
| Control 4 | | | 164 | 63.08 ± 1.89 | 12 |
| Control 5 | | | 144 | 66.93 ± 3.59 | 6 |
| Control 6 | | | 135 | 64.61 ± 2.57 | 6 |
| Control 7 | | | 122 | 66.11 ± 5.57 | 7 |

**Supplementary Statistical analysis**

Fig. 6A: Body Collapses. Statistically significant scores indicated in red.

**Pooled data**

| Z test results | | | |
|---|---|---|---|
| | | Z score | p Value |
| Control | TeTX | -7.0147 | <0.00001 |

**Pairwise Comparisons**

| Z test results | | | |
|---|---|---|---|
| | | Z score | p Value |
| Control1 | Control2 | 0.6395 | 0.52218 |
| Control1 | Control3 | 1.6064 | 0.1704 |
| Control1 | Control4 | 1.0451 | 0.29372 |
| Control2 | Control3 | 0 | 1 |
| Control2 | Control4 | 0 | 1 |
| Control3 | Control4 | 0 | 1 |
| Control1 | TeTX1 | -0.6932 | 0.4902 |
| Control1 | TeTX2 | -2.8791 | 0.00398 |
| Control1 | TeTX3 | -2.7099 | 0.00672 |
| Control1 | TeTX4 | -3.2439 | 0.0012 |
| Control1 | TeTX5 | -3.7566 | 0.00016 |
| Control2 | TeTX1 | -0.9321 | 0.35238 |
| Control2 | TeTX2 | -2.1564 | 0.03078 |
| Control2 | TeTX3 | -2.0468 | 0.04036 |
| Control2 | TeTX4 | -2.39 | 0.01684 |
| Control2 | TeTX5 | -2.694 | 0.00714 |
| Control3 | TeTX1 | -2.3323 | 0.0198 |
| Control3 | TeTX2 | -5.1786 | <0.00001 |
| Control3 | TeTX3 | -4.9411 | <0.00001 |
| Control3 | TeTX4 | -5.6775 | <0.00001 |
| Control3 | TeTX5 | -6.3364 | <0.00001 |
| Control4 | TeTX1 | -1.5212 | 0.12852 |
| Control4 | TeTX2 | -3.4432 | 0.00058 |
| Control4 | TeTX3 | -3.2787 | 0.00104 |
| Control4 | TeTX4 | -3.7928 | 0.00016 |
| Control4 | TeTX5 | -4.266 | <0.00001 |

### Fig. 6B: Statistical analysis
### Analysis of Range (Knee Height)

#### Quantiles

| Level | Minimum | 10% | 25% | Median | 75% | 90% | Maximum |
|---|---|---|---|---|---|---|---|
| Control | 1.356529 | 1.605726 | 1.732048 | 1.883241 | 2.196335 | 2.512926 | 3.01324 |
| TeXT | 1.739098 | 2.260857 | 2.532573 | 3.146695 | 3.385301 | 4.151452 | 5.048257 |

#### Means and Std Deviations

| Level | Number | Mean | Std Dev | Std Err Mean | Lower 95% | Upper 95% |
|---|---|---|---|---|---|---|
| Control | 504 | 1.9810563 | 0.33479 | 0.0149127 | 1.9517574 | 2.0103553 |
| TeXT | 630 | 3.1088092 | 0.7461462 | 0.0297272 | 3.0504327 | 3.1671857 |

#### Tests that the Variances are Equal

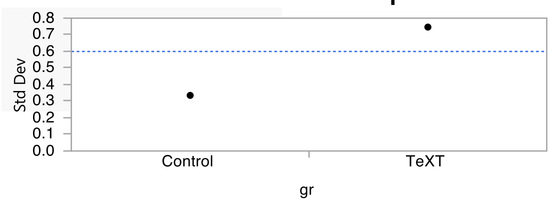


| Level | Count | Std Dev | MeanAbsDif to Mean | MeanAbsDif to Median |
|---|---|---|---|---|
| Control | 504 | 0.3347900 | 0.2772036 | 0.2696084 |
| TeXT | 630 | 0.7461462 | 0.5714468 | 0.5700685 |

| Test | F Ratio | DFNum | DFDen | p-Value |
|---|---|---|---|---|
| O'Brien[.5] | 132.3435 | 1 | 1132 | <.0001* |
| Brown-Forsythe | 167.4121 | 1 | 1132 | <.0001* |
| Levene | 169.2823 | 1 | 1132 | <.0001* |
| Bartlett | 309.7565 | 1 | . | <.0001* |
| F Test 2-sided | 4.9671 | 629 | 503 | <.0001* |

#### Welch's Test

Welch Anova testing Means Equal, allowing Std Devs Not Equal

| F Ratio | DFNum | DFDen | Prob > F |
|---|---|---|---|
| 1149.8353 | 1 | 913.1 | <.0001* |

| t Test |
|---|
| 33.9092 |

### Fig. 6C: Statistical analysis
### Analysis of Range (TMP Angle)

| WATSON-WILLIAMS F-TESTS | | | | | |
|---|---|---|---|---|---|
| Variables (& observations) | F | p | df | df2 | Est. Mean |
| Controls & TeTX (504 & 630) | 430.895 | < 1E-12 | 1 | 1132 | 62.188 |

Table SS1: Force Test

| BASIC STATISTICS | | | | | | | | | | |
|---|---|---|---|---|---|---|---|---|---|---|
| Variable | Control5 | Control6 | Control7 | Control8 | Control9 | TeTX1 | TeTX2 | TeTX3 | TeTX4 | TeTX5 |
| Data Type | Angles | Angles | Angles | Angles | Angles | Angles | Angles | Angles | Angles | Angles |
| Number of Observations | 6 | 6 | 7 | 16 | 11 | 3 | 3 | 5 | 5 | 3 |
| Mean Vector (µ) | 66.928 | 64.619 | 66.12 | 64.214 | 63.081 | 68.41 | 63.643 | 68.453 | 69.297 | 66.444 |
| Circular Standard Deviation | 3.592 | 2.57 | 5.569 | 1.997 | 1.89 | 2.291 | 1.516 | 2.997 | 3.114 | 3.069 |
| One Sample Tests | | | | | | | | | | |
| Rayleigh Test (Z) | 5.976 | 5.988 | 6.934 | 15.981 | 10.988 | 2.995 | 2.998 | 4.986 | 4.985 | 2.991 |
| Rayleigh Test (p) | < 1E-12 | < 1E-12 | < 1E-12 | 1.97E-07 | 1.3E-06 | 0.034 | 0.033 | 0.001 | 0.001 | 0.034 |

**Watson-Williams F test (control vs TeTX chickens)**

| Source | d.f. | SS | MS | F | P-Value |
|---|---|---|---|---|---|
| Columns | 1 | 0.921084 | 0.921084 | 1.722648 | 0.225754 |
| Residual | 8 | 7.749183 | 0.968648 | | |
| Total | 9 | 8.670267 | | | |

Table SS2: Comparison of maximum delta per stride

| ANOVA results | | | | | |
|---|---|---|---|---|---|
| Source | SS | df | MS | F | prob>F |
| Groups | 220.94 | 6 | 36.8233 | 17.58 | 1.05E-18 |
| Error | 1181.62 | 564 | 2.0951 | | |
| Total | 1402.56 | 570 | | | |

### Quantiles

| Level | Minimum | 10% | 25% | Median | 75% | 90% | Maximum |
|---|---|---|---|---|---|---|---|
| Control3 | 1.367596 | 2.843941 | 3.388987 | 4.047114 | 4.895785 | 5.652417 | 6.379741 |
| Control4 | 1.18938 | 2.550268 | 3.253339 | 4.307534 | 5.311383 | 5.995616 | 7.939445 |
| TeTX1 | 0.34284 | 3.260252 | 3.817822 | 4.999565 | 6.136319 | 7.413171 | 11.8446 |
| TeTX2 | 2.176949 | 3.825371 | 4.514134 | 5.429668 | 5.755859 | 6.651419 | 10.36888 |
| TeTX3 | 1.613126 | 2.939709 | 4.042566 | 4.794431 | 4.957137 | 5.403267 | 6.341758 |
| TeTX4 | 2.95143 | 3.469008 | 4.127639 | 5.111682 | 5.50772 | 5.921419 | 9.004741 |
| TeTX5 | 1.642539 | 4.121106 | 4.789288 | 5.538087 | 6.422167 | 9.005017 | 10.24161 |

### Means and Std Deviations

| Level | Number | Mean | Std Dev | Std Err Mean | Lower 95% | Upper 95% |
|---|---|---|---|---|---|---|
| Control3 | 137 | 4.1500897 | 1.0749626 | 0.0918402 | 3.96847 | 4.3317093 |
| Control4 | 115 | 4.3271494 | 1.321059 | 0.1231894 | 4.0831122 | 4.5711866 |
| TeTX1 | 97 | 5.1120102 | 1.8971602 | 0.1926274 | 4.7296477 | 5.4943726 |
| TeTX2 | 36 | 5.3208716 | 1.3849789 | 0.2308298 | 4.8522622 | 5.7894811 |
| TeTX3 | 27 | 4.5069349 | 1.0170163 | 0.1957249 | 4.1046166 | 4.9092531 |
| TeTX4 | 49 | 4.9071678 | 1.164942 | 0.1664203 | 4.5725574 | 5.2417782 |
| TeTX5 | 110 | 5.8255975 | 1.7156012 | 0.1635762 | 5.5013949 | 6.1498002 |

### Means Comparisons

### Comparisons with a control using Dunnett's Method

Control Group = Control1

### Confidence Quantile

| \|d\| | Alpha |
|---|---|
| 2.61488 | 0.05 |

### LSD Threshold Matrix

| Level | Abs(Dif)-LSD | p-Value |
|---|---|---|
| TeTX5 | 1.191 | <.0001* |
| TeTX2 | 0.462 | 0.0001* |
| TeTX1 | 0.46 | <.0001* |
| TeTX4 | 0.127 | 0.0102* |
| TeTX3 | -0.44 | 0.7654 |
| Control3 | -0.3 | 0.8836 |
| Control4 | -0.46 | 1.0000 |

Positive values show pairs of means that are significantly different.

Fig. S8A: Phase analysis

| BASIC STATISTICS | | | | | | | |
|---|---|---|---|---|---|---|---|
| Variable | GFP1 | GFP2 | TeTX1 | TeTX2 | TeTX3 | TeTX4 | TeTX5 |
| Data Type | Angles | Angles | Angles | Angles | Angles | Angles | Angles |
| Number of Observations | 75 | 31 | 204 | 155 | 114 | 79 | 142 |
| Mean Vector (μ) | 182.369 | 182.384 | 184.674 | 182.293 | 180.784 | 180.502 | 181.97 |
| Length of Mean Vector (r) | 0.827 | 0.897 | 0.847 | 0.856 | 0.863 | 0.818 | 0.823 |
| Circular Standard Deviation | 35.366 | 26.708 | 33.003 | 32.01 | 31.064 | 36.291 | 35.787 |
| One Sample Tests | | | | | | | |
| Rayleigh Test (Z) | 51.239 | 24.946 | 146.4 | 113.443 | 84.966 | 52.892 | 96.131 |
| Rayleigh Test (p) | < 1E-12 | 7.34E-11 | < 1E-12 | < 1E-12 | < 1E-12 | < 1E-12 | < 1E-12 |

| WATSON-WILLIAMS F-TESTS | | | | | |
|---|---|---|---|---|---|
| Variables (& observations) | F | p | df | df2 | Est. Mean |
| Multi-sample test using: | | | | | |
| GFP1 (75) | | | | | |
| GFP2 (31) | | | | | |
| TeTX1 (204) | | | | | |
| TeTX2 (155) | | | | | |
| TeTX3 (114) | | | | | |
| TeTX4 (79) | | | | | |
| TeTX5 (142) | 0.25 | 0.959 | 6 | 793 | 182.466 |

Figure S8B – Ranges of knee height

### Quantiles

| Level | Minimum | 10% | 25% | Median | 75% | 90% | Maximum |
|---|---|---|---|---|---|---|---|
| Control_GFP1 | 1.583211 | 1.642068 | 1.732058 | 1.853918 | 2.036332 | 2.302025 | 2.338401 |
| Control_GFP2 | 1.356529 | 1.505224 | 1.553307 | 1.852974 | 2.061026 | 2.171198 | 2.220186 |
| Control_NoElec1 | 2.151395 | 2.170454 | 2.196808 | 2.381171 | 2.644546 | 2.742689 | 3.01324 |
| Control_NoElec2 | 1.574461 | 1.606336 | 1.661235 | 1.781316 | 1.840663 | 1.870193 | 1.914784 |
| TeTX1 | 2.221573 | 2.267482 | 2.648965 | 3.163871 | 3.358293 | 3.798085 | 3.948563 |
| TeTX2 | 3.180134 | 3.230696 | 3.278528 | 3.375239 | 3.61459 | 4.067007 | 4.314338 |
| TeTX3 | 2.190001 | 2.274949 | 2.343899 | 2.461626 | 2.703694 | 3.196665 | 3.322023 |
| TeTX4 | 1.739098 | 1.764574 | 1.977728 | 2.614597 | 2.978234 | 3.317682 | 3.538927 |
| TeTX5 | 2.771483 | 2.882091 | 3.133971 | 3.380367 | 4.820164 | 5.003136 | 5.048257 |

### Oneway Anova
### Summary of Fit

| | |
|---|---|
| Rsquare | 0.732832 |
| Adj Rsquare | 0.730932 |
| Root Mean Square Error | 0.425585 |
| Mean of Response | 2.607586 |
| Observations (or Sum Wgts) | 1134 |

### Analysis of Variance

| Source | DF | Sum of Squares | Mean Square | F Ratio | Prob > F |
|---|---|---|---|---|---|
| Label | 8 | 558.91272 | 69.8641 | 385.7283 | <.0001* |
| Error | 1125 | 203.76287 | 0.1811 | | |
| C. Total | 1133 | 762.67560 | | | |

### Means for Oneway Anova

| Level | Number | Mean | Std Error | Lower 95% | Upper 95% |
|---|---|---|---|---|---|
| Control_GFP1 | 126 | 1.91458 | 0.03791 | 1.8402 | 1.9890 |
| Control_GFP2 | 126 | 1.83644 | 0.03791 | 1.7620 | 1.9108 |
| Control_NoElec1 | 126 | 2.41935 | 0.03791 | 2.3450 | 2.4937 |
| Control_NoElec2 | 126 | 1.75386 | 0.03791 | 1.6795 | 1.8282 |
| TeTX1 | 126 | 3.05139 | 0.03791 | 2.9770 | 3.1258 |
| TeTX2 | 126 | 3.50734 | 0.03791 | 3.4330 | 3.5817 |
| TeTX3 | 126 | 2.57487 | 0.03791 | 2.5005 | 2.6493 |
| TeTX4 | 126 | 2.54391 | 0.03791 | 2.4695 | 2.6183 |
| TeTX5 | 126 | 3.86654 | 0.03791 | 3.7921 | 3.9409 |

Figure S8C – Ranges of TMP joint

| BASIC STATISTICS | | | | | | | | | |
|---|---|---|---|---|---|---|---|---|---|
| Variable | Control1 | Control2 | Control3 | Control4 | TeTX1 | TeTX2 | TeTX3 | TeTX4 | TeTX5 |
| Data Type | Angles | Angles | Angles | Angles | Angles | Angles | Angles | Angles | Angles |
| Number of Observations | 126 | 126 | 126 | 126 | 126 | 126 | 126 | 126 | 126 |
| Mean Vector (μ) | 56.804 | 41.427 | 54.864 | 44.118 | 82.279 | 71.793 | 64.171 | 72.488 | 72.866 |
| Circular Standard Deviation | 16.583 | 18.673 | 9.687 | 12.587 | 22.21 | 25.234 | 21.269 | 17.213 | 12.024 |

| WATSON-WILLIAMS F-TESTS | | | | | |
|---|---|---|---|---|---|
| Variables (& observations) | F | p | df | df2 | Est. Mean |
| Multi-sample test using: | | | | | |
| Control1 (126) | | | | | |
| Control2 (126) | | | | | |
| Control3 (126) | | | | | |
| Control4 (126) | | | | | |
| TeTX1 (126) | | | | | |
| TeTX2 (126) | | | | | |
| TeTX3 (126) | | | | | |
| TeTX4 (126) | | | | | |
| TeTX5 (126) | 76.168 | < 1E-12 | 8 | 1125 | 62.188 |

Fig. S1

A

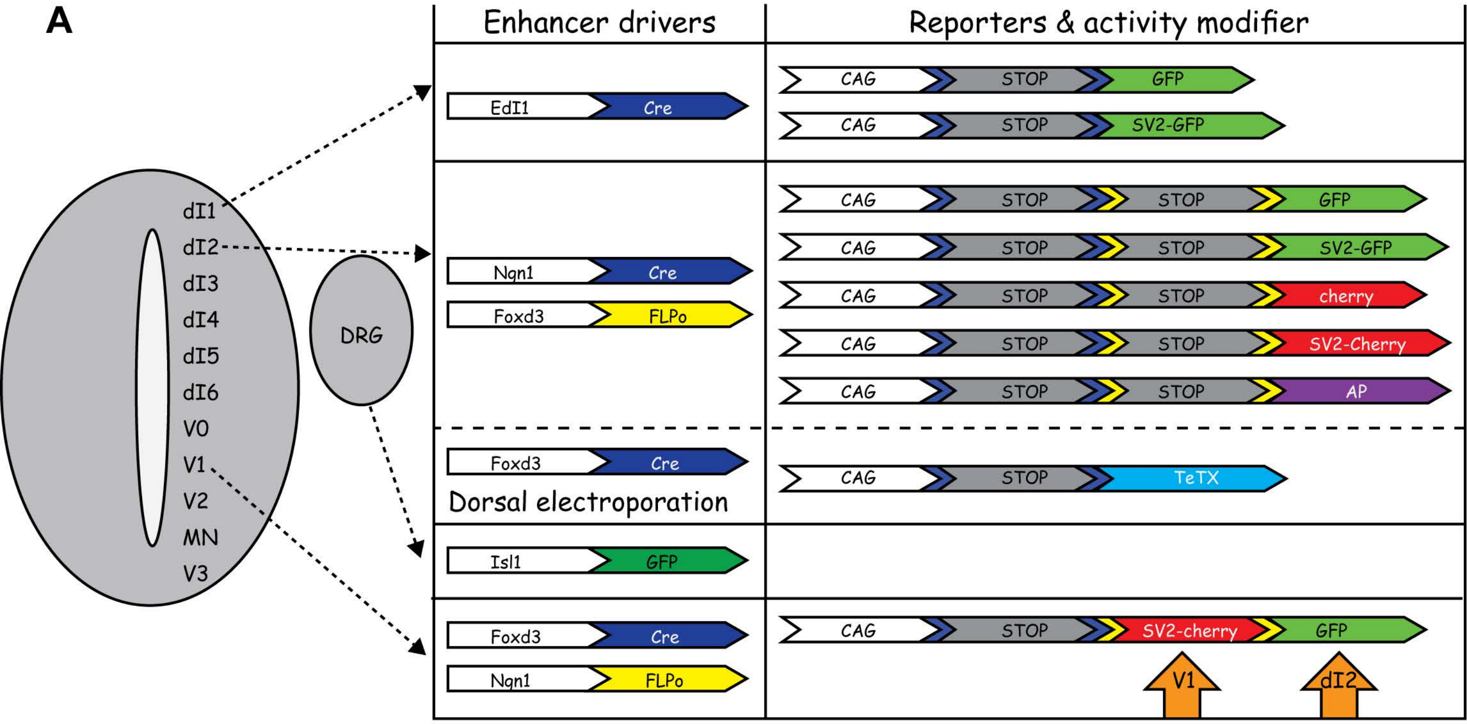


B

| | Axonal projection | Neurotransmitter | Soma location | Pre-MN |
|---|---|---|---|---|
| dI1c | C | E | V | No |
| dI1i | I | E | M | Yes |
| dI2 | C | E | M/V | No |
| dI3 | I | E | D/M | Yes |
| dI4 | I | I | D | No |
| dIL | I | E/I | D | |
| dI5 | I | E | D | Yes |
| dI6 | I/C | I | M/V | Yes |
| V0 | C | I/E | M/V | Yes |
| V1 | I | I | M/V | Yes |
| V2 | I | E/I | M/V | Yes |
| V3 | C | E | D | Yes |

Fig. S2: Differential expression of TFs in dI2 INs.

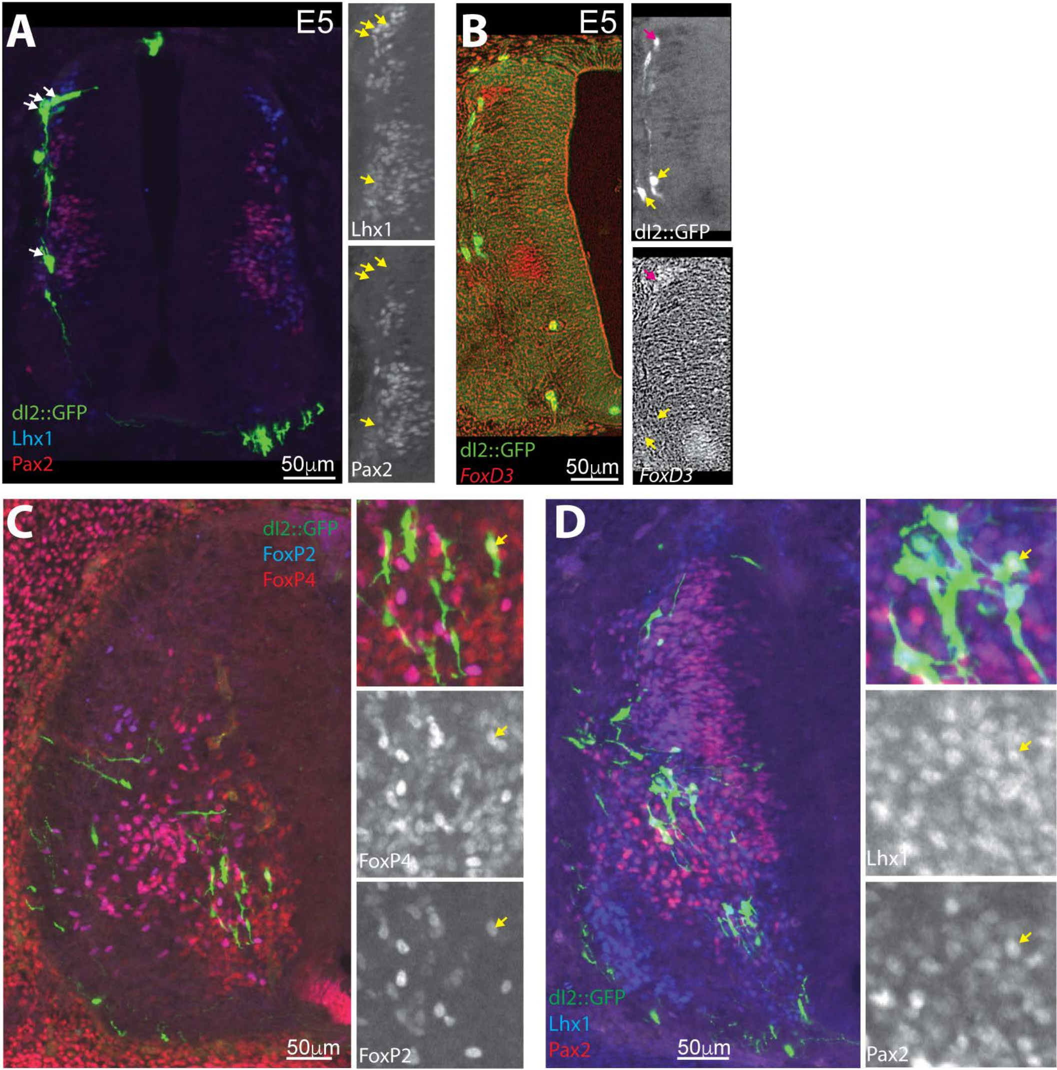

Fig. S3: dI2s Neuro-transmitter phenotype and soma localization.

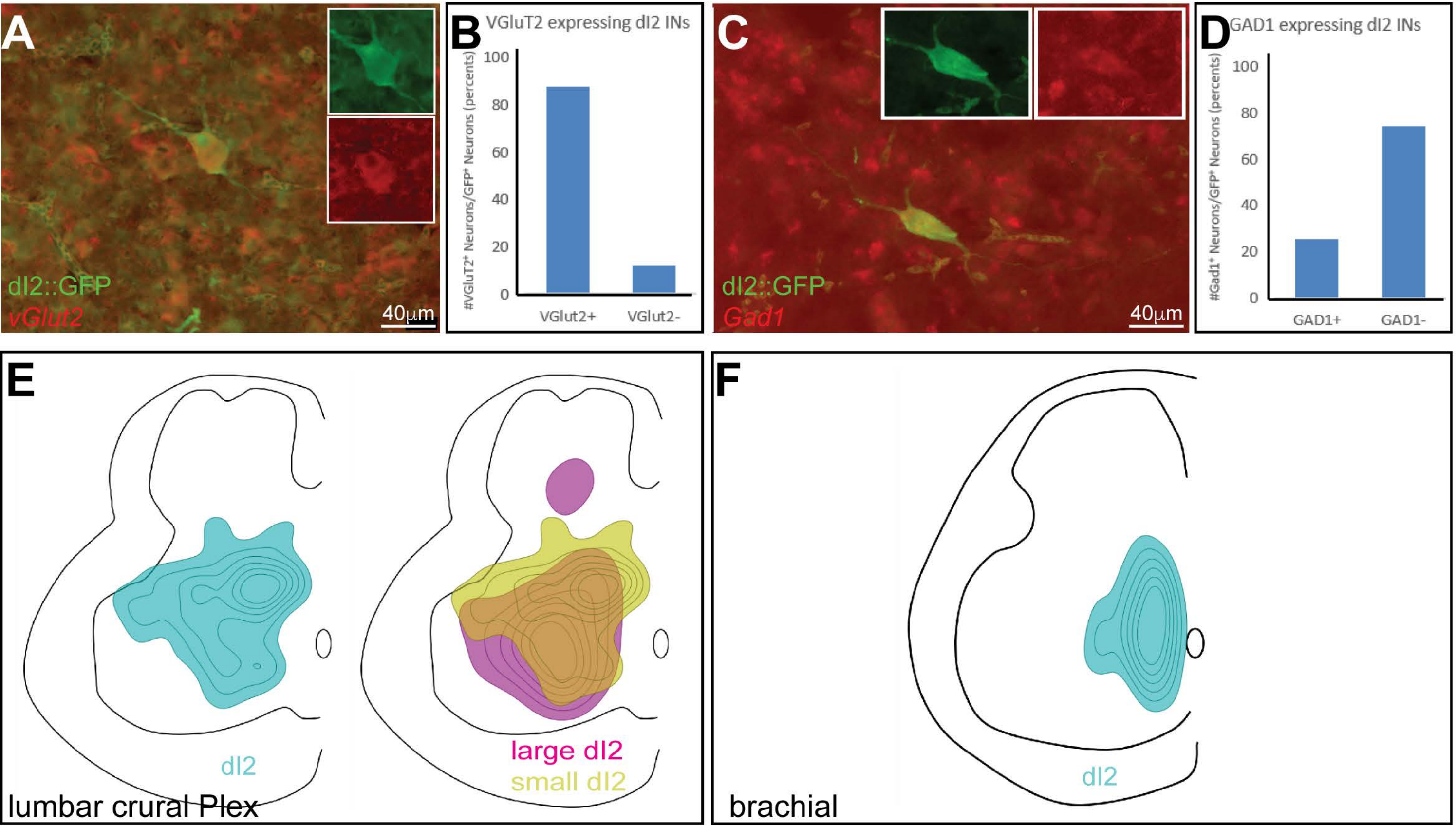

Fig. S4: dI2s densities at the crural plexus level, innervation of deep cerebellar nuclei and projection to the cerebellum

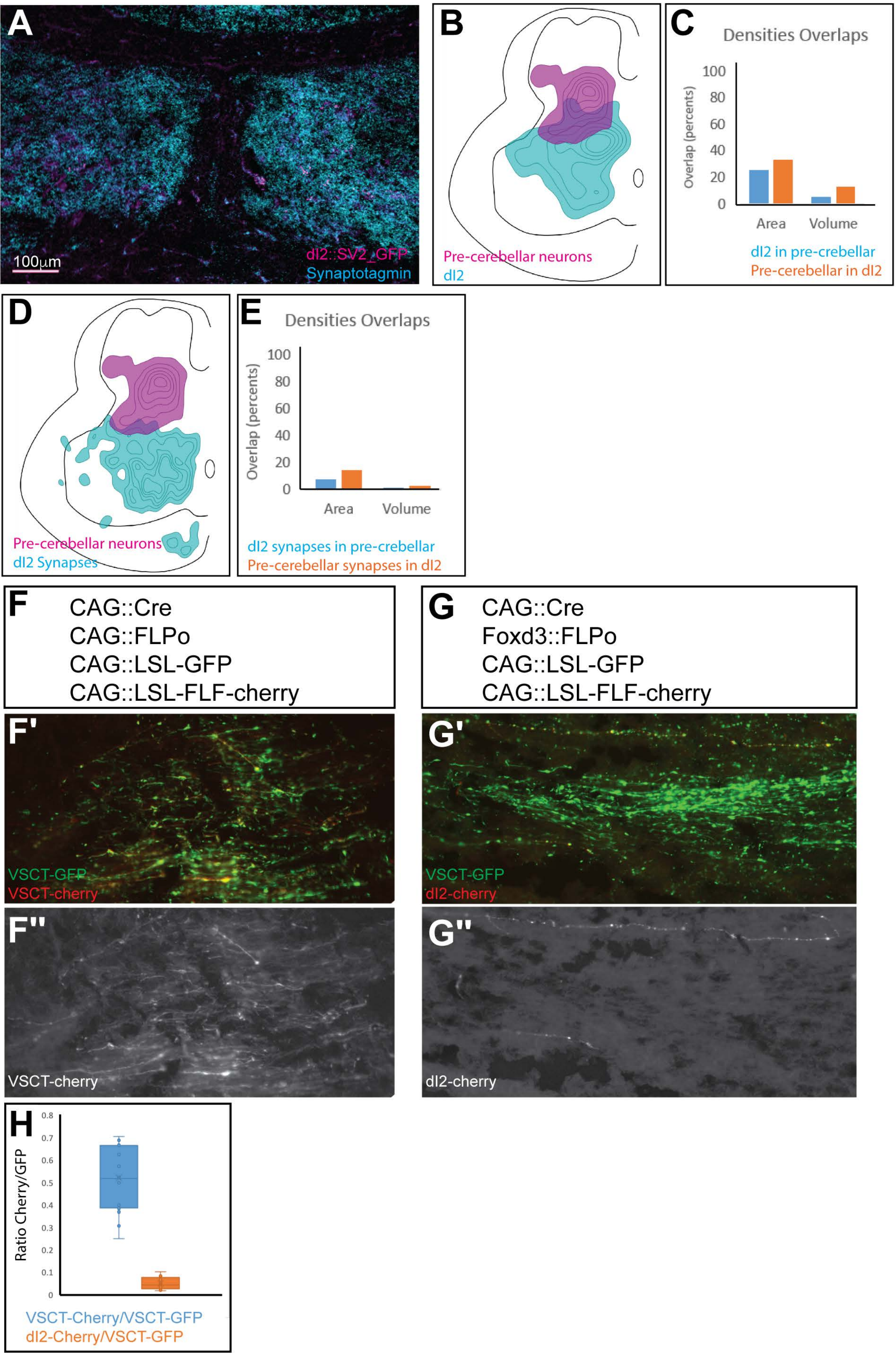

Fig. S5: dI1i are pre-MNs

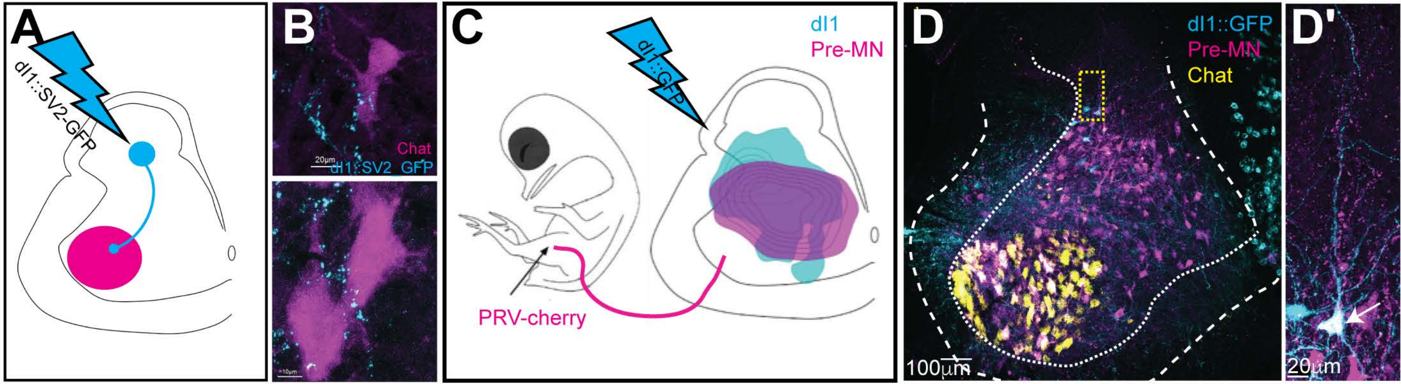

Fig. S6: Input of 5HT and dI1 to dI2 neurons at the level of the crural plexus

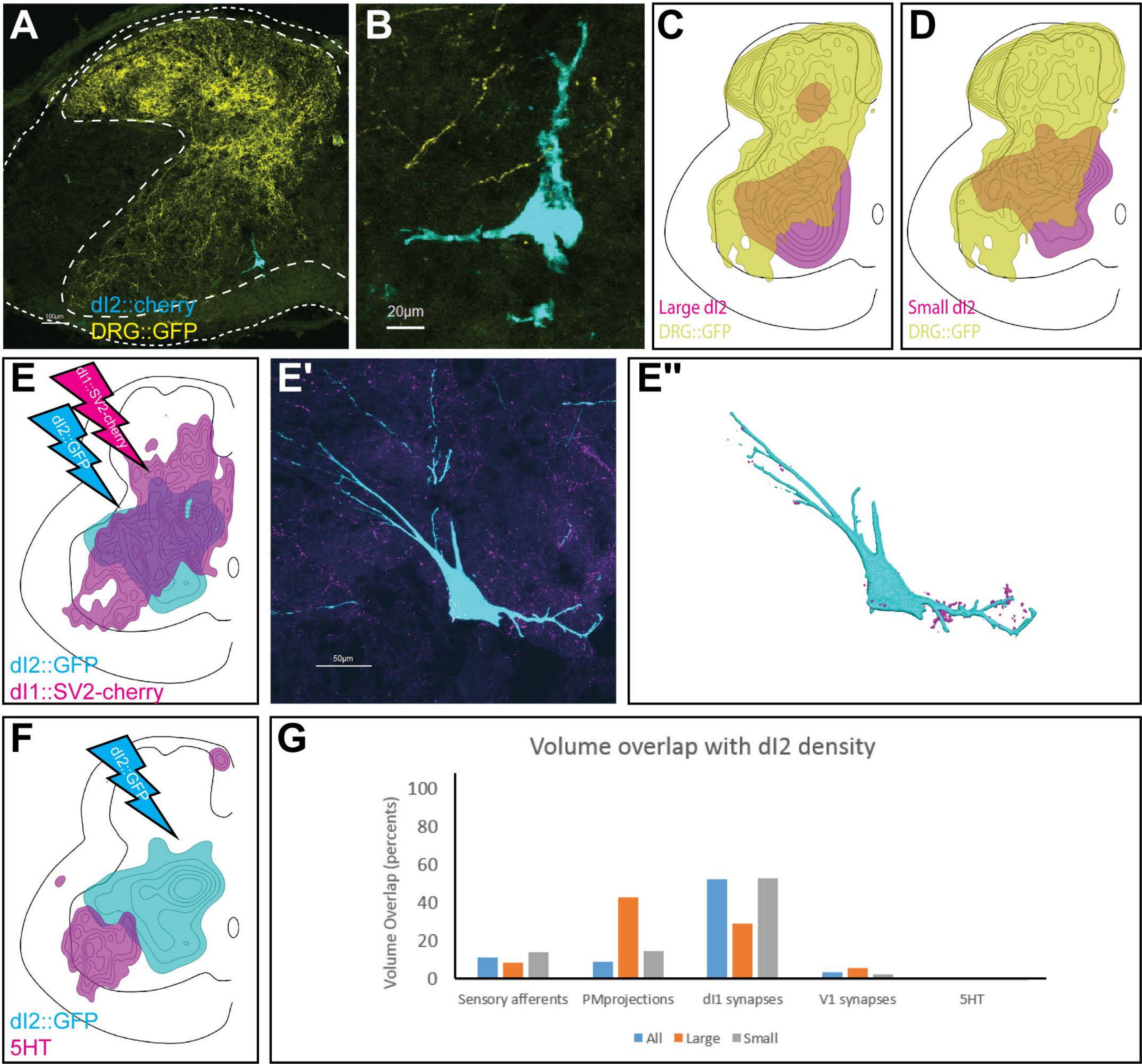

Fig. S7: Spinal targets of dI2

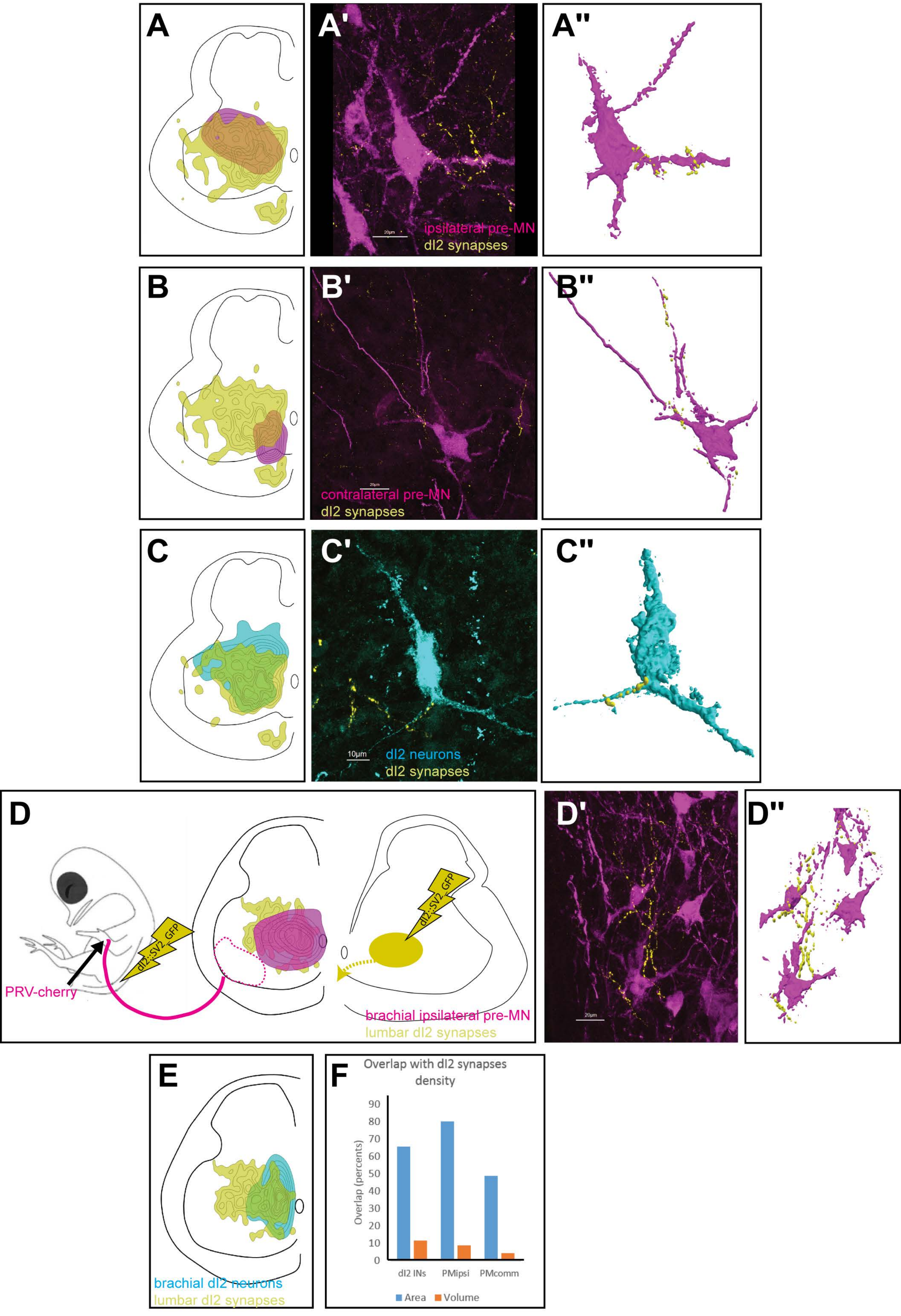

Fig. S8: Locomotion characteristics of control and TeTX-treated chicks.

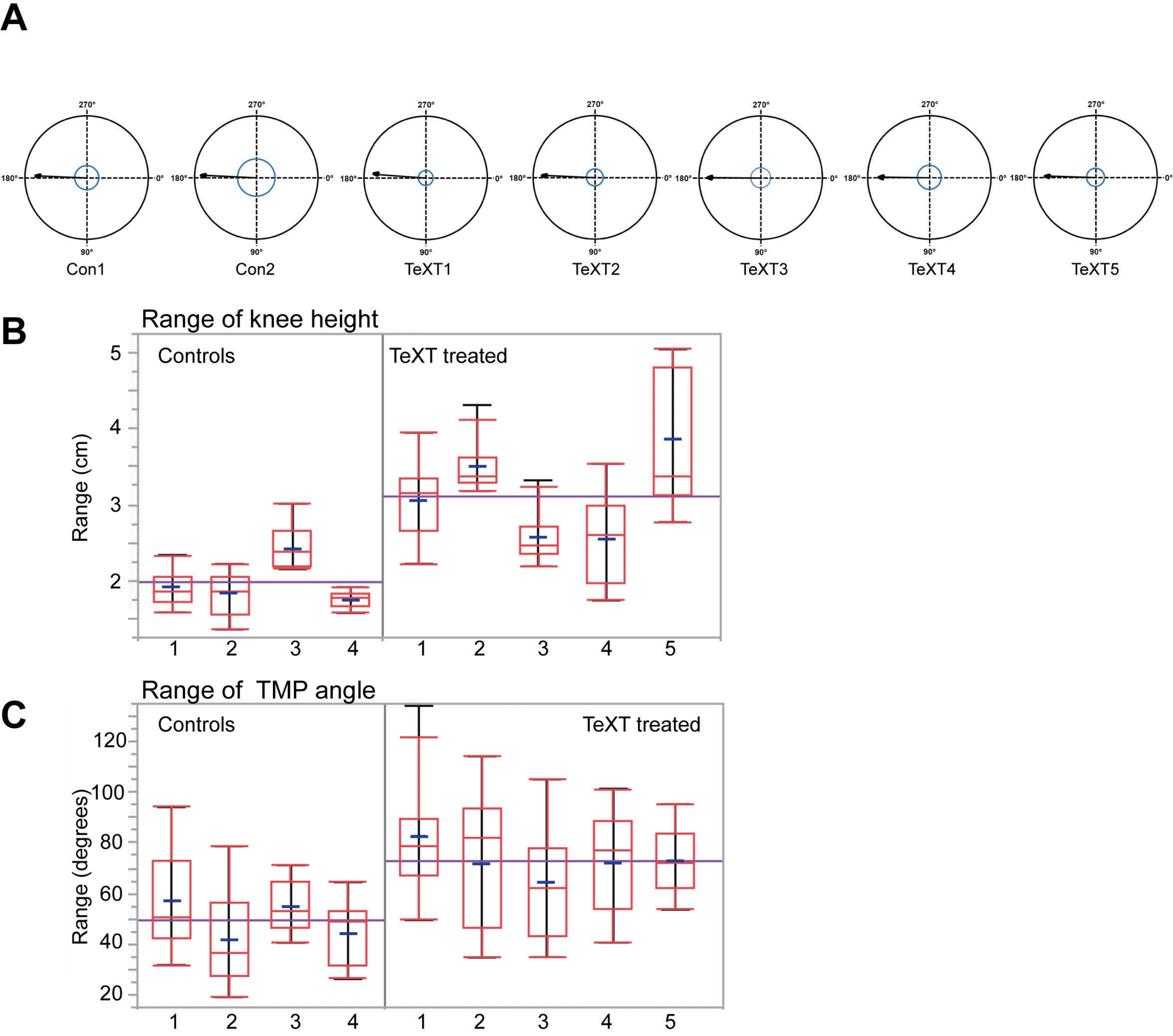
